# Water status shapes responses to ultraviolet radiation in two stress-tolerant mosses

**DOI:** 10.1101/2022.09.14.508022

**Authors:** Jenna T. B. Ekwealor, Suzanne Kosina, Anderson T. Silva, Benjamin Bowen, Trent Northen, Melvin J. Oliver, Brent D. Mishler

## Abstract

Mosses cycle between hydrated, metabolically active periods and desiccated quiescence, with water status constraining when they can actively respond to stressors such as ultraviolet radiation (UVR). We tested how UVR responses depended on water status in the closely related mosses *Syntrichia ruralis* and *S. caninervis*. Using a fully factorial chronic UVR × hydration experiment, we combined transcriptomics with untargeted metabolomics of soluble and cell-wall-bound fractions. Across both species, drying elicited the broadest transcriptomic and soluble metabolomic responses. The breadth of the UVR-associated transcriptomic response diverged between species across hydration states: the number of transcripts differing between UVR treatments increased during drying in *S. ruralis* but decreased in *S. caninervis*. Wall-associated chemistry showed a strong species-specific UVR signature, with nearly all significant interaction features attributable to species × UVR, while UVR × hydration interactions were rare. Nevertheless, conditional UVR responses in the wall-associated metabolome were much broader in hydrated than drying tissue. Water status therefore altered the extent of UVR-associated changes in transcript and metabolite abundance, and the two species differed in how those responses were distributed across hydration states and molecular compartments.

## 1 INTRODUCTION

Mosses experience repeated transitions between hydrated, metabolically active periods and desiccated quiescence. Because they are poikilohydric, water availability determines when active responses to environmental stress are possible: as tissues dry, metabolic activity progressively declines and eventually stops (Bewley, 1979; Alpert and Oliver, 2002; Proctor et al., 2007). Desiccation tolerance requires surviving water loss and the desiccated state and restoring function upon rehydration, and can involve constitutive protection, responses induced during drying, and repair of damage accumulated during drying or while tissues are desiccated (Oliver et al., 2005; Proctor et al., 2007; Oliver et al., 2020). Environmental exposures such as solar radiation, however, continue through drying and desiccation, leaving mosses exposed to ultraviolet radiation (UVR) even as their capacity for active response and repair declines. Responses initiated while tissues are hydrated or early in drying may therefore be especially important if their effects persist into desiccation.

Water status may alter not only the capacity to respond to UVR, but also the conditions under which UVR is experienced by the tissue. As mosses dry, changes in leaf orientation and tissue geometry can alter the exposure of photosynthetic tissues to incoming radiation (Seel et al., 1992). For example, photon absorption in dry *Syntrichia ruraliformis* was 50–60% lower than in hydrated plants because of leaf curling (Seel et al., 1992). Hydration state can also change the consequences of UVR exposure itself: desiccated tissues of two Antarctic mosses accumulated substantially less UV-B-induced DNA damage than hydrated tissues, whereas desiccation was not protective in a third species (Turnbull et al., 2009). At the same time, declining cellular water changes molecular mobility and the physicochemical environment in which cellular reactions occur (Oliver et al., 2020; Farrant and Hilhorst, 2021). UVR encountered by hydrated and drying tissues may therefore have different consequences and elicit different molecular responses.

Wall-associated chemistry is especially relevant to this timing problem because UVR-screening compounds at or near the tissue surface are positioned to intercept incoming radiation and can remain in place as tissues dry (Clarke and Robinson, 2008). UVR-absorbing phenolics, including flavonoids, contribute to bryophyte photoprotection, and cell-wall-bound compounds can constitute a substantial fraction of the total UVR-absorbing pool (Taipale and Huttunen, 2002; Clarke and Robinson, 2008; Robinson and Waterman, 2014; Waterman et al., 2017). This investment may also carry a cost: across bryophytes, greater concentrations of wall-bound UVR-absorbing compounds are associated with lower maximum photosynthetic capacity, consistent with reduced CO_2_ diffusion through phenolic-rich walls (Perera Castro et al., 2026; Ekwealor and Simonin, 2026). Wall-associated UVR protection may therefore be especially consequential in poikilohydric plants, where carbon gain is already restricted to finite periods of hydration.

The moss genus *Syntrichia* is an ecologically diverse lineage that includes several highly desiccation-tolerant dry-land species (Mishler, 2007; Jauregui-Lazo et al., 2023). *Syntrichia caninervis* is characteristic of arid and semiarid habitats, including biological soil crusts, whereas *S. ruralis* occupies a much broader range of habitats and moisture regimes (Mishler, 2007). The two species also differ physiologically: comparative experiments ranked *S. caninervis* as more desiccation tolerant than *S. ruralis*, although tolerance also varied substantially among populations (Oliver et al., 1993). In *S. caninervis*, dark pigmentation varies with light environment and UVR exposure, and long-term reduction of natural UVR alters pigment composition and photosynthetic physiology (Ekwealor, 2020; Ekwealor and Fisher, 2020; Ekwealor et al., 2021). These ecological and physiological contrasts provide a tractable comparison of how closely related poikilohydric plants respond to UVR across changing water status.

Here, we used a fully factorial chronic UVR × hydration experiment to test whether and how UVR responses change across the hydration–drying cycle in *S. caninervis* and *S. ruralis*. We combined transcriptomics with untargeted metabolomics of methanol-soluble and cell-wall-bound fractions to ask three questions: how broadly do drying and UVR alter transcript and metabolite abundance, how does hydration state modify the UVR response, and how do those responses differ between species and among the transcriptome, soluble metabolome, and cell-wall-bound metabolome? Based on extensive physiological and molecular reorganization during desiccation in bryophytes and other resurrection plants, we predicted broad changes in transcript and metabolite abundance during drying (Oliver, 1991; Phillips et al., 2002; Liu et al., 2018; Oliver et al., 2020). *Syntrichia caninervis* is strongly associated with arid biocrust habitats where prolonged summer drought limits periods of physiological activity, whereas *S. ruralis* occupies a broader range of habitats and moisture regimes (Mishler, 2007; Clark et al., 2025). Previous acute-exposure work showed that hydrated *S. caninervis* can mount a substantial transcriptomic response to UVR, including responses of early light-inducible proteins (Ekwealor and Mishler, 2021), but did not test how that response changes as tissues dry. Based on their contrasting relationships between UVR exposure and water availability, we predicted that hydration state would modify UVR responses differently in the two species. We expected the UVR response of *S. ruralis* to be strongest while tissues were hydrated, when plants can actively respond to radiation, whereas the UVR response of *S. caninervis* would vary less between hydrated and drying tissue, consistent with greater reliance on constitutive or persistent protection during prolonged dry periods. Given the potential for wall-associated compounds to persist as tissues dry, we also expected the two species to differ in their wall-associated responses to UVR.

## 2 MATERIALS AND METHODS

### 2.1 Experimental conditions

To compare the effects of UVR and hydration state on the metabolome and transcriptome of two *Syntrichia* species, we grew clonal isolates of *S. caninervis* and *S. ruralis* under controlled growth-chamber conditions. We cultivated shoots from a previously isolated clone of *S. caninervis* collected in southern Nevada, USA (Stark NV-107; Clark County, New-berry Mountains, Christmas Tree Pass; UNLV), and an isolated clone of *S. ruralis* collected near the Bow River in Calgary, Canada (Brinda 9108; UNLV), from fragments on 1.2% agar containing an inorganic nutrient solution (Hoagland and Arnon, 1950). We grew cultures in Magenta GA-7 plant culture boxes under an 18-h photoperiod at 16 °C during the light period and 8 °C during the dark period.

We placed culture boxes under eight T8 reptile bulbs (ReptiSun 10.0 UVB, Zoo Med Laboratories Inc., San Luis Obispo, CA, USA). We manipulated UV exposure using acrylic windows placed over the culture boxes. We covered sixteen cultures of each species with 7.6 × 7.6 cm, 3.175-mm-thick UVR-filtering acrylic windows (OP-3 acrylic, Acrylite, Sanford, ME, USA), sealed with Parafilm. We wrapped the sides of each box in aluminum foil so that incident light passed through the acrylic window. UVR-filtering windows transmitted approximately 90% of visible radiation but approached 0% transmittance between 425 and 400 nm. UVR-transmitting controls used otherwise comparable Polycast Solacryl SUVT acrylic (Spartech, Maryland Heights, MO, USA), which transmitted at least 90% of visible and UVR-A/B radiation before declining to near 0% between 275 and 250 nm. Both window types transmitted approximately 90% of photosynthetically active radiation (PAR; 400–700 nm) and filtered UVR-C.

We positioned lamps approximately 10 cm above the agar surface and 3 cm above the acrylic windows. The high-UVR treatment had a UVR-A/B flux of 36 µmol m_−2_ s_−1_ and a UV-B fluence rate of 0.15 mW cm_−2_; the low-UVR treatment had a UVR-A/B flux of 5 µmol m_−2_ s_−1_ and no detectable UV-B (0.00 mW cm_−2_). Both treatments received approximately 120 µmol m_−2_ s_−1_ PAR. We measured PAR and UVR-A/B using a LightScout Sensor Reader with LightScout Quantum and UV sensors (Spectrum Technologies, Aurora, IL, USA), collecting measurements at a minimum of 30 positions throughout the chamber and three measurements per position. We measured UV-B fluence at several locations using a handheld radiometer (SKU 430 Apollo Display Meter, Skye Instruments Ltd., Llandrindod

Wells, UK) with the UV-B sensor covered by the same acrylic used for the experimental treatments. We grew cultures for three months and randomized box positions at least every other week. We monitored relative humidity with an iButton Hygrochron (Maxim Integrated, San Jose, CA, USA; mean chamber RH = 80%).

### 2.2 Hydration treatments and tissue collection

To test the combined effects of UVR and drying, we collected tissues either while hydrated or following a slow-drying treatment. We collected seven biological replicates for each species × UVR × hydration combination, using separate biological replicates for transcriptomics and metabolomics (Table S1). We made all collections at least 1 h into the light period. We clipped above-ground tissue at the base to minimize contamination from the agar growth medium.

For transcriptomics, we placed three biological replicates per treatment combination in 1.6-mL microcentrifuge tubes with a small hole in the cap, immediately froze them in liquid nitrogen, and stored them at -80 °C. For metabolomic we placed four separate biological replicates per treatment combination in open microcentrifuge tubes and lyophilized them for 48 h using a Labconco FreeZone 2.5-L Benchtop Freeze Dry System with a Fisherbrand Maxima C Plus M6C Vacuum Pump before storage at -80 °C.

For the drying treatment, we placed clipped tissue on filter paper in culture dishes, covered it with the same UVR-filtering or UVR-transmitting acrylic used during cultivation, and returned it to the growth chamber. We dried tissues at approximately 80% RH (-29.54 MPa) until severe dehydration was evident from leaf curling and other morphological signs associated with approaching desiccation. We selected these conditions to produce slow drying while maintaining exposure to the experimental UVR treatment and growth temperature, and to reduce the likelihood that transcript sequestration associated with more rapid drying would dominate changes in transcript abundance (Wood and Oliver, 1999; Stark et al., 2017; Slate et al., 2018). We collected drying-treated samples after five days for transcriptomics and six days for metabolomics, then either froze them in liquid nitrogen and stored them at -80 °C or lyophilized and stored them at -80 °C, respectively.

## 3 METABOLOMICS

### 3.1 Metabolite extraction

We performed separate extractions targeting methanol-soluble and cell-wall-bound metabolites, following previously published protocols (Semerdjieva et al., 2003; Clarke and Robinson, 2008; Waterman et al., 2017, 2018). For the methanol-soluble fraction, we pulverized 20–50 mg lyophilized tissue in a bead mill, resuspended it in water, froze it at -80 °C, and pulverized it again. We added 1 mL methanol and sonicated samples for 1 h at room temperature in the dark. After centrifugation, we collected the supernatant and repeated the methanol extraction three additional times using 0.5 mL methanol, pooling the supernatants across extractions. We dried the pooled extracts, resuspended them in methanol containing internal standard while keeping samples cold, vortexed and sonicated them in ice water, centrifuged them, and passed the supernatant through 0.22-µm filters before LC–MS analysis.

For the cell-wall-bound fraction, we sequentially washed the residual pellets with 1 M sodium chloride, methanol, 1:1 methanol:chloroform, and acetone and then dried them. We hydrolyzed the washed pellets overnight in 1 M sodium hydroxide in the dark, collected the supernatant, acidified it with 1 M hydrochloric acid, and lyophilized it. We resuspended the resulting material in ice-cold LC–MS-grade methanol, vortexed and sonicated it in an ice bath, centrifuged it, and dried the recovered supernatant. We then resuspended the dried extract in methanol containing internal standard, centrifuged and filtered it through 0.22-µm filters, and transferred the filtrate to LC–MS vials.

### 3.2 Untargeted LC-MS profiling and molecular networking

We used untargeted LC–MS/MS to profile metabolite features separately from the soluble and cell-wall-bound fractions. We analyzed methanol-soluble extracts using hydrophilic interaction liquid chromatography (HILIC) and cell-wall-bound extracts using reversed-phase C18 chromatography. We acquired positive- and negative-ionization data and processed the two ionization modes separately.

We processed LC–MS/MS data using MZmine 2.39 (Pluskal et al., 2010). We detected centroided ions separately in MS1 and MS2 scans using noise thresholds of 10,000 and 100, respectively. We constructed chromatograms using the ADAP chromatogram builder (Myers et al., 2017), requiring at least five scans per group, a group-intensity threshold of 200,000, a minimum highest intensity of 500,000, and an m/z tolerance of 0.002 Da or 10 ppm. We deconvoluted chromatograms using the ADAP wavelets algorithm with a signal-to-noise threshold of 10 and paired MS2 scans to features within 0.025 Da and 0.2 min. We grouped isotopic features using an m/z tolerance of 20 ppm and retention-time tolerance of 0.1 min, allowing charge states up to 3 and using the most intense isotope as the representative isotope. We aligned features across samples using the join aligner with an m/z tolerance of 20 ppm and retention-time tolerance of 0.2 min, with equal weighting of m/z and retention time. We retained aligned rows represented by at least two samples, filtered duplicate features using an m/z tolerance of 0.0001 Da or 10 ppm and a retention-time tolerance of 0.2 min, and gap-filled the resulting feature table using an intensity tolerance of 5%, an m/z tolerance of 0.001 Da or 10 ppm, and a retention-time tolerance of 0.2 min. MZmine exported both peak-height and peak-area feature tables; we used peak height as the feature-abundance measure for downstream analyses.

We performed molecular networking using the feature-based molecular networking workflow in the Global Natural Products Social Molecular Networking platform (GNPS) (Wang et al., 2016). For GNPS submission, we exported MS/MS spectra only for features associated with an MS2 scan. We removed MS/MS fragment ions within ±17 Da of the precursor and window-filtered spectra by retaining the six most intense fragment ions within each ±50-Da window. We set both precursor- and fragment-ion mass tolerances to 0.05 Da. We required network edges to have a cosine similarity score >0.70 and more than six matched peaks and retained edges only when each node occurred among the other node’s ten most similar spectra. We limited molecular family size to 100 nodes by removing lower-scoring edges as necessary. We also searched spectra against GNPS libraries using the same filtering criteria and retained matches at cosine scores ≥ 0.70 with at least six matched peaks. We treated library matches as putative annotations rather than confirmed metabolite identities.

We additionally used targeted m/z–retention-time information to support putative annotation of selected untargeted soluble features. We matched manually accepted targeted compound-ion candidates to untargeted MZmine features in the corresponding ionization mode using target-specific m/z tolerances and retention-time windows, retaining match-quality and ambiguity information for each candidate. Targeted data were used only as annotation evidence and were not used as the abundance matrix for differential-abundance analyses. All effect estimates, adjusted P values, and significance calls were calculated from the untargeted peak-height feature tables. Where a feature had both a unique targeted m/z–retention-time annotation and a GNPS library annotation, the targeted annotation was used as the preferred display label. When targeted and GNPS annotations agreed, evidence from both sources was retained; when they conflicted, the feature was flagged as having conflicting candidate identities. Features associated with multiple targeted compound names were retained as ambiguous targeted-evidence sets; when no independent unambiguous annotation was available, no compound identity was assigned. Because authentic-standard co-analysis under the same LC–MS conditions was not available, unique targeted or GNPS compound annotations were treated as putative MSI level 2 annotations, whereas unresolved ambiguous or conflicting annotations were not assigned a compound identity. Annotation source, match quality, and supporting evidence are reported in Table S9.

For molecular-family abundance profiles, we standardized each feature separately across all biological samples in both species, UVR treatments, and hydration states using the mean and sample standard deviation of its log-normalized abundance. We calculated these feature-level z-scores after excluding missing values from the mean and standard-deviation calculations and assigned a standardized value of zero to features with zero or invalid standard deviation. Family-level lines and points represent the median standardized abundance of member features within each species × UVR × hydration group.

### 3.3 Untargeted metabolomic quality control and feature filtering

We processed untargeted datasets separately for soluble negative-ion, soluble positive-ion, cell-wall-bound negative-ion, and cell-wall-bound positive-ion analyses. We excluded features detected in fewer than two biological samples. We also removed features that showed strong evidence of association with extraction controls, treatment controls, or other blanks. We classified a feature as control-associated when it was detected in at least 50% of the relevant controls and the ratio of maximum biological abundance to maximum control abundance was < 3. We also excluded features with direct contaminant annotations from either GNPS or targeted annotation sources from the biological analyses.

For each analytical run, we normalized retained peak heights to total sample signal to reduce between-sample variation in overall ion abundance. We scaled the resulting relative abundances by 10^6^ and applied a log(1 + *x*) transformation to reduce right skew and improve variance stability for PCA and linear-model analyses. We used these normalized, transformed abundances for principal component analyses (Fig. S1). For exploratory joint PCA of each metabolomic fraction, we matched samples across positive- and negative-ionization modes. Within each ionization mode, we removed zero-variance features and centered and scaled each retained feature across samples. To balance the contributions of the two ionization modes, we divided each mode-specific feature matrix by the square root of its number of retained features, concatenated the two matrices, and performed PCA without additional centering or scaling. This weighting gave the positive- and negative-ionization blocks approximately equal aggregate variance in the joint ordination. Ellipses in the joint PCA plots are 95% normal-theory summaries and were used only for visualization. We excluded one sample (sample 52) from both cell-wall-bound datasets because it was a strong multivariate outlier in both ionization modes, while the corresponding biological replicate behaved normally in the soluble datasets, consistent with a fraction-specific analytical or extraction problem (Fig. S2; Table S1).

### 3.4 Differential metabolite abundance

We analyzed differential abundance separately for each of the four metabolomic datasets using linear models implemented in limma (Ritchie et al., 2015). For each feature, we fit a full factorial model including species, UVR treatment, hydration state, and all two- and three-way interactions:

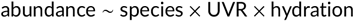

We coded factors using sum-to-zero contrasts so that each factorial main effect represented an average effect across the other factors rather than a comparison restricted to a single reference treatment. We used empirical Bayes moderation of residual variances with trend adjustment and robust estimation.

In addition to factorial term tests, we defined planned contrasts to quantify average species, UVR, and hydration effects; species × UVR, species × hydration, UVR × hydration, and three-way interactions; species-specific UVR and hydration effects; within-species UVR × hydration interactions; hydration responses within each UVR treatment; and UVR responses within each hydration state. We applied Benjamini–Hochberg correction separately within each factorial term or planned contrast and considered features with adjusted *P* values < 0.05 significant.

For multivariate inference, we performed PERMANOVA on Euclidean distances calculated from the same total-signal-normalized abundance matrices used for PCA using adonis2 in the R package *vegan* (Anderson, 2001; Oksanen et al., 2026). We fit full factorial models including species, UVR, hydration state, and their interactions, with 9,999 permutations. Because sequential sums of squares can depend on term order in unbalanced designs, we retained alternate factor orders as a sensitivity check. We assessed multivariate dispersion among treatment groups using bias-adjusted distances to group centroids with betadisper and evaluated differences by permutation using permutest in *vegan* (Anderson, 2001; Oksanen et al., 2026). Dispersion results are reported in Table S10.

We evaluated robustness to alternative normalization methods, control-associated feature thresholds, inclusion of cell-wall-bound sample 52, linear adjustment for injection order, and conservative exclusion of all members of molecular families containing a known contaminant annotation (Fig. S5; Table S11).

## 4 TRANSCRIPTOMICS

### 4.1 RNA extraction and sequencing

We extracted total RNA from frozen tissue using the RNeasy Plant Mini Kit (Qiagen, Germantown, MD, USA) according to the manufacturer’s protocol. We removed residual DNA using the TURBO DNA-free Kit (Invitrogen, Waltham, MA, USA). We assessed RNA quantity, purity, and integrity using a Qubit fluorometer and Bioanalyzer 2100. We prepared libraries using the NEBNext Ultra II RNA Library Prep with Sample Purification Beads kit and sequenced them as 50-bp single-end reads on an Illumina HiSeq 4000 at the QB3 Vincent J. Coates Genomics Sequencing Laboratory.

### 4.2 Read processing, alignment, and counting

We processed raw sequencing reads separately by species. We cleaned technical sequencing lanes from the same biological sample separately and then merged them after preprocessing. We filtered reads using fastp (Chen et al., 2018) with a minimum Phred quality score of 20 and a minimum retained length of 30 bp and used automatic adapter detection. We assessed cleaned reads with FastQC (Andrews, 2010).

We aligned reads from each species, *S. caninervis* and *S. ruralis*, to its corresponding published reference genome (Zhang et al., 2024; Silva et al., 2021) using STAR (Dobin et al., 2013). Alignments allowed up to 20 multimapping locations and a maximum mismatch proportion of 0.04 relative to read length. We generated gene-level counts from exon alignments with featureCounts (Liao et al., 2014) using gene_id as the grouping attribute. Functional annotations and gene descriptions were taken from the annotations accompanying the published *S. caninervis* and *S. ruralis* reference genomes (Silva et al., 2021; Zhang et al., 2024). Because the libraries were single-end and unstranded, we counted reads as unstranded single reads. Before differential-abundance analysis, we excluded one *S. caninervis* high-UVR hydrated library because its assigned gene-count total (46,125 reads across both sequencing lanes) was below 10% of the *S. caninervis* median library size (Table S1).

### 4.3 Differential transcript abundance

We analyzed differential transcript abundance (gene-level RNA-seq abundance quantified from reads assigned to annotated genes) separately for *S. ruralis* and *S. caninervis* using DESeq2 (Love et al., 2014) because we aligned each species to its own reference genome. Within each species, the model was:

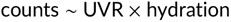

We retained genes when they had at least 10 reads in at least as many samples as the smallest UVR × hydration treatment cell for that species. We then applied DESeq2 default size-factor normalization and dispersion estimation. For each species, we tested the average effect of UVR across hydration states, the average effect of drying across UVR treatments, the UVR × hydration interaction, UVR effects separately within hydrated and dry tissues, and hydration effects separately within low- and high-UVR treatments. We defined mean effects as equal-weight averages across the two levels of the interacting factor. We adjusted *P* values using the Benjamini–Hochberg procedure and considered transcripts with adjusted *P* values < 0.05 significantly differentially abundant.

We used variance-stabilizing transformation for PCA and sample-distance visualization. Because we analyzed transcriptomes against species-specific reference genomes, we restricted direct cross-species comparisons of individual genes to explicitly identified orthologous genes rather than treating reference gene IDs as directly equivalent between species.

### 4.4 Transcript-response modules

We identified within-species transcript-response modules from the same four conditional treatment responses used in the differential-abundance analysis: the UVR effect while hydrated, the UVR effect during drying, the drying effect under low UVR, and the drying effect under high UVR. We used the existing modeled log_2_ fold-change estimates for these contrasts and identified modules separately for each species.

For each species, we column-standardized the four-contrast response profiles and performed Ward.D2 hierarchical clustering on Euclidean distances. We evaluated dendrogram cuts from *k* = 3 to 12 and selected the cut maximizing the median transcript-to-cluster-centroid Pearson correlation plus 0.01 log(*k*), among cuts in which every provisional cluster contained at least 30 transcripts. This procedure selected *k* = 12 for both species.

Within each provisional cluster, we then refined membership iteratively using the unscaled four-contrast response profiles. Transcripts were retained when their response profile correlated with the module centroid at *r* ≥ 0.90 and matched the centroid in sign for all four contrasts. After each filtering step, the centroid was recalculated and membership was reevaluated, for up to 30 iterations; transcripts failing these criteria were left unassigned, and refined clusters containing fewer than 30 transcripts were not retained.

We next considered refined modules for iterative post hoc merging. Candidate module pairs were required to have Pearson correlation ≥ 0.95 between their row-standardized four-condition centroids, identical signs for all four response contrasts, and Euclidean distance below a species-specific cutoff defined by the largest gap in the observed pairwise centroid-distance distribution (0.297 for *S. caninervis* and 0.099 for *S. ruralis*). At each iteration, the closest eligible pair was merged, and transcript membership was reevaluated using the same *r* ≥ 0.90 correlation, sign-matching, and minimum-size criteria. Merging continued until no eligible module pairs remained.

For the primary analysis, we retained merged modules containing at least one transcript significant for the UVR × hydration interaction at Benjamini–Hochberg-adjusted *P* < 0.05. The four conditional UVR and drying contrasts were used to characterize the response structure of these retained modules but did not determine module retention. For visualization, we used variance-stabilized transcript abundances averaged within each of the four treatment combinations and standardized each transcript across those four conditions. The standardized values were used only to display response shape and were not used to redefine module membership.

### 4.5 Cross-species ortholog comparison

We identified one-to-one orthologs between the two species using OrthoFinder v3.1.5 with the DendroBLAST workflow (Emms and Kelly, 2019). Protein sequences were generated from the published *S. caninervis* and *S. ruralis* genome annotations, retaining the longest translated isoform per gene. OrthoFinder used DIAMOND for sequence searches, FAMSA for multiple-sequence alignment, and FastTree for gene-tree inference. We retained orthogroups containing exactly one gene from each species, yielding 9,044 one-to-one ortholog pairs. We reconciled ortholog identifiers with the gene-level DESeq2 feature identifiers using the reference annotations used for RNA-seq analysis. Where the *S. ruralis* orthology output contained transcript-level identifiers, we mapped them to genes using explicit parent relationships in the GFF annotation and retained only unique, unambiguous mappings to DESeq2-tested genes. Of the 9,044 one-to-one ortholog pairs, 8,935 were present in both reference annotations, and 7,003 pairs (77.4%) mapped unambiguously to tested genes in both species (Table S2).

For descriptive comparisons of species-specific responses, we classified mapped orthologs for each planned contrast as significant in both species in the same direction, significant in both in opposite directions, significant only in *S. ruralis*, significant only in *S. caninervis*, or not significant in either species. We defined significance independently within each species and contrast at Benjamini–Hochberg-adjusted *P* < 0.05. We used fold-change direction to distinguish same-from opposite-direction responses only when the ortholog was significant in both species.

For direct cross-species inference, we constructed a joint count matrix containing the 7,003 mapped one-to-one orthologs and samples from both species and fit a DESeq2 model with species, UVR, water status, and all interactions:

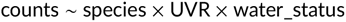

We used unshrunk DESeq2 Wald estimates and their corresponding standard errors. We tested species differences in the UVR effect separately within hydrated and drying tissues using contrasts from this joint model and applied Benjamini–Hochberg correction separately within each contrast. Of the 7,003 mapped orthologs, 6,999 had non-missing adjusted P-values for these contrasts; four were untestable after model filtering. We also used variance-stabilized counts from the 7,003-ortholog joint matrix for the cross-species transcriptomic PCA.

## 5 RESULTS

### 5.1 Global molecular structure and breadth of treatment responses

Species identity and water status structured all three molecular datasets (Fig. 1). In the joint transcriptomic PCA of 7,003 one-to-one orthologs, species separated primarily along PC1, which explained 72.94% of the variance, while water status separated samples within species along PC2, which explained 12.67% (Fig. 1A). Joint, mode-balanced PCAs of the soluble and cell-wall-bound metabolomes likewise separated the species while retaining clear within-species structure associated with hydration state and UVR treatment (Fig. 1B,C); separate transcriptomic and ionization-mode PCA views are shown in Fig. S1.

**FIGURE 1.**
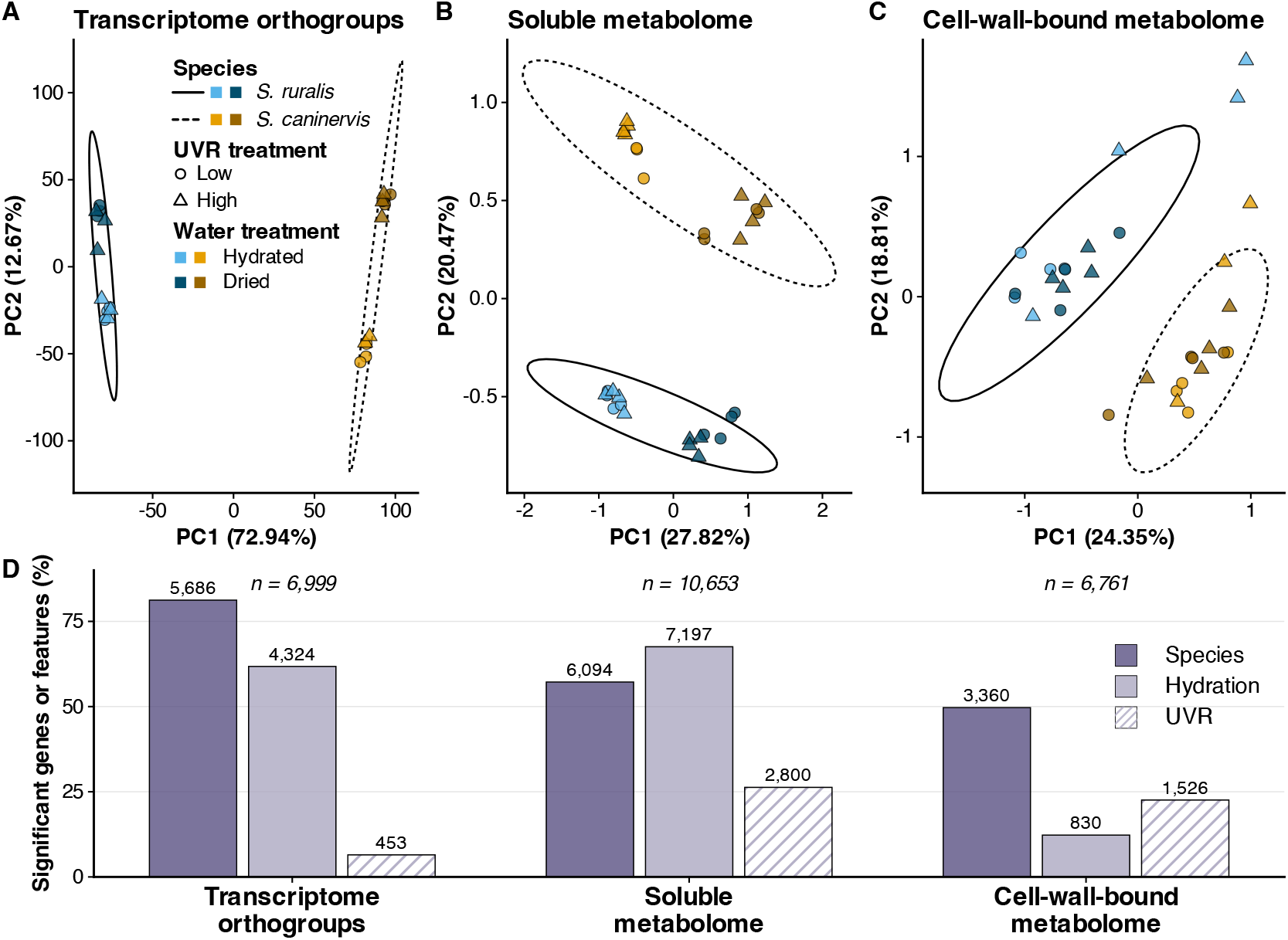
Global molecular structure and breadth of environmental responses across omics layers. (A) Joint orthogroup transcriptomic PCA based on variance-stabilized transcript abundance for 7,003 mapped one-to-one orthologs. (B) Mode-balanced joint PCA of the methanol-soluble metabolome integrating separately processed positive- and negative-ionization features. (C) Mode-balanced joint PCA of the cell-wall-bound metabolome integrating separately processed positive- and negative-ionization features. In panels A–C, point encodings indicate species, UVR treatment, and water status as shown in the legends. (D) Percentage of tested orthogroups or metabolomic features significant for species, hydration, and UVR main effects at Benjamini–Hochberg-adjusted P < 0.05, with raw significant counts annotated and the number of tested genes or features shown for each molecular compartment. Transcriptomic effects in panel D are from the joint orthogroup DESeq2 model, whereas metabolomic effects are from the limma factorial models; positive- and negative-ionization datasets were analyzed separately for inference and pooled only for this visual summary.

The breadth of significant treatment responses differed among molecular compartments (Fig. 1D). Species identity affected 5,686 of 6,999 testable orthogroups, 6,094 of 10,653 soluble-metabolomic features, and 3,360 of 6,761 cell-wall-bound features (Fig. 1D). Across the joint orthogroup transcriptome, hydration affected 4,324 orthogroups compared with 453 for UVR. Across the separately analyzed ionization modes of the soluble metabolome, hydra-tion affected 7,197 features compared with 2,800 for UVR. In the cell-wall-bound metabolome, UVR affected 1,526 features compared with 830 affected by hydration across ionization modes. Species identity was a major source of variation across datasets, while the relative effects of water status and UVR differed by molecular compartment: water status affected more transcripts and soluble features, whereas UVR affected more wall-associated features.

### 5.2 Hydration state reverses the relative transcriptomic UVR response of the two species

Drying affected many more transcripts than the average UVR effect across hydration states in both species, but the UVR response depended on water status. Within *S. ruralis*, transcripts from 8,889 of 21,892 tested genes were significantly differentially abundant for the average drying effect across UVR treatments, while transcripts from 738 genes were differentially abundant for the average UVR effect across hydration states. Within *S. caninervis*, transcripts from 8,228 of 13,963 tested genes were differentially abundant for the average drying effect across UVR treatments, while transcripts from 1,011 genes were differentially abundant for the average UVR effect across hydration states.

Hydration state changed the breadth of the UVR response in opposite directions in the two species (Fig. 3A). In *S. ruralis*, transcripts from only 179 genes were significantly differentially abundant between high and low UVR in hydrated tissue, whereas transcripts from 875 genes were differentially abundant during drying. In contrast, *S. caninervis* showed a larger UVR response while hydrated: transcripts from 791 genes were differentially abundant between high and low UVR in hydrated tissue compared with transcripts from 295 genes during drying. UVR × hydration interactions were significant for 130 genes in *S. ruralis* and 64 genes in *S. caninervis*.

The leading annotated UVR × hydration transcripts also differed between species (Fig. 2). In *S. caninervis*, an early light-inducible protein (ELIP) transcript showed the strongest UVR × hydration interaction; its abundance increased strongly under high UVR while hydrated, whereas transcript abundance was already high during drying and showed little additional UVR response. Other prominent annotated interaction transcripts in *S. caninervis* included a ferric-iron-binding protein and *NRT2.2*. In *S. ruralis*, the strongest annotated interaction responses included a YLS9-like across the four conditional treatment contrasts. (C) Corresponding retained *S. ruralis* modules. For panels B and C, standardized transcript abundance is shown across low and high UVR separately for hydrated and drying tissue; thick lines show module centroids, ribbons show the interquartile range, and faint lines show sampled module members. Module labels report total membership and the number of transcripts significant for the UVR × hydration interaction. Only modules containing at least one UVR × hydration-interaction-significant transcript are shown. gene, an exostosin-family gene, and a member of the iron/ascorbate-dependent oxidoreductase family, each showing different UVR responses between hydrated and drying tissue.

**FIGURE 2.**
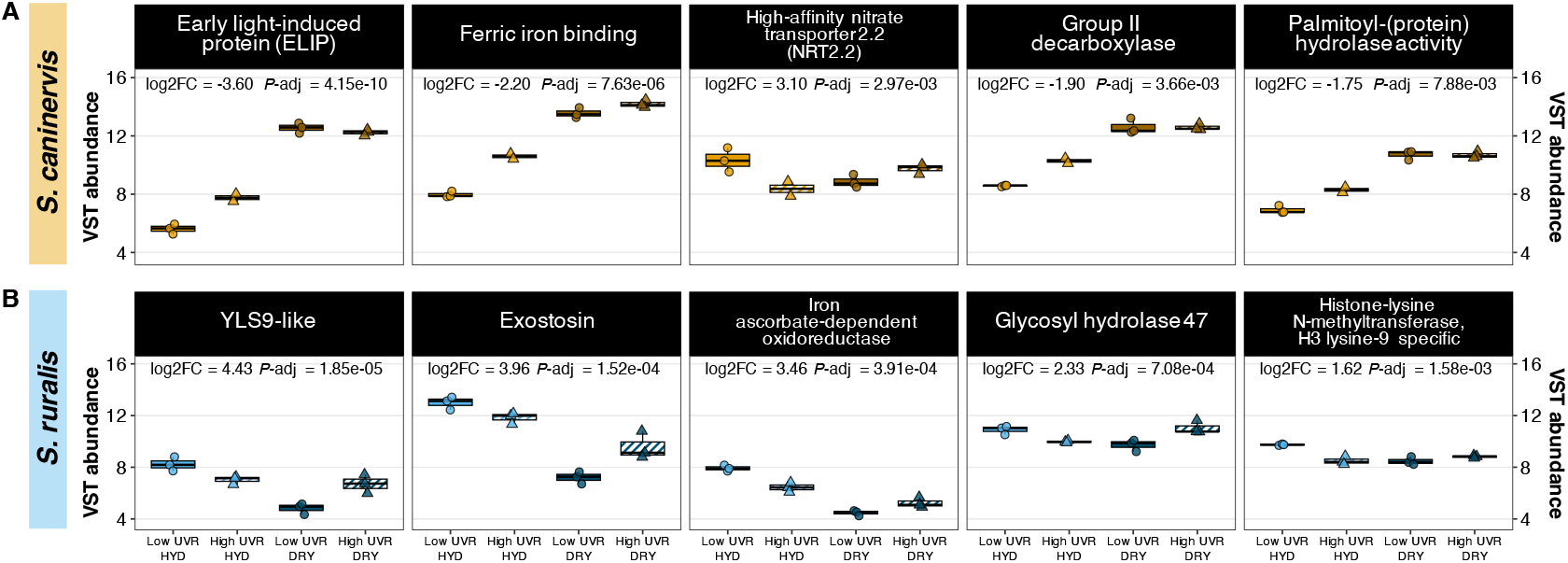
Leading UVR × hydration transcript responses differ between the two Syntrichia species. (A) The five strongest annotated UVR × hydration interaction transcripts in S. caninervis. (B) The corresponding five strongest annotated interaction transcripts in S. ruralis. For each transcript, variance-stabilized transcript abundance is shown across low and high UVR in hydrated (HYD) and drying (DRY) tissue, with biological replicates overlaid on boxplots. Displayed statistics are the within-species UVR × hydration interaction log2 fold change and Benjamini–Hochberg-adjusted P value. Transcripts were ranked by adjusted P value among annotated genes in the within-species UVR × hydration interaction test.

These opposite responses also appeared in transcript-response modules (Fig. 3). Of nine merged modules in *S. caninervis*, four contained at least one transcript significant for the UVR × hydration interaction, whereas two of seven merged modules in *S. ruralis* met this criterion. The retained *S. caninervis* modules contained 124, 74, 62, and 212 transcripts, respectively, whereas the two retained *S. ruralis* modules contained 208 and 291 transcripts. The ELIP transcript belonged to *S. caninervis* Module 3, a 74-transcript module containing three transcripts significant for the UVR × hydration interaction; its response vector correlated strongly with the module centroid (*r* = 0.981). Retained-module membership and summary statistics are reported in Tables S3 and S4, and effect patterns and significance structure are shown in Fig. S3.

**FIGURE 3.**
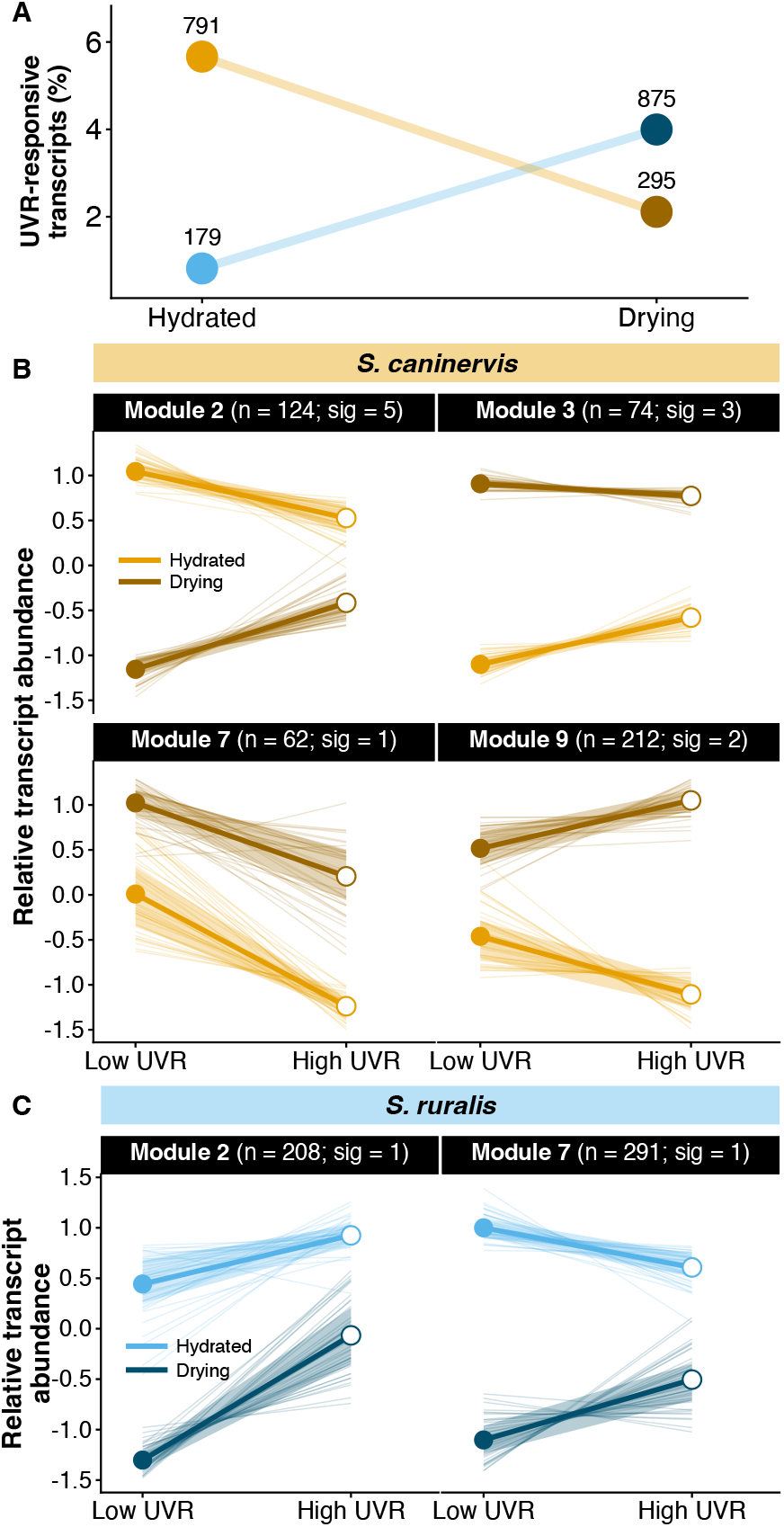
Hydration state structures both the breadth and module-level patterns of UVR-associated transcript responses. (A) Percentage of tested transcripts differing significantly between high and low UVR within hydrated and drying tissue in each species, with significant-transcript counts shown for each contrast (S. ruralis: 179 hydrated and 875 drying; S. caninervis: 791 hydrated and 295 drying). (B) Retained S. caninervis transcript-response modules, each containing at least one transcript significant for the UVR × hydration interaction at Benjamini–Hochberg-adjusted P < 0.05; module membership required correlation ≥ 0.90 to the module centroid and complete sign agreement

Drying responses remained extensive under both UVR treatments in both species, although the number of responsive genes also varied with UVR environment. In *S. ruralis*, 7,101 genes differed between dry and hydrated tissue under low UVR and 4,334 under high UVR. In *S. caninervis*, the corresponding values were 7,234 and 5,707 genes.

### 5.3 Drying responses are broadly shared, whereas UVR responses diverge between species

The cross-species ortholog analysis separated broadly shared drying responses from more species-specific UVR responses. Of 9,044 one-to-one ortholog pairs, 7,003 mapped unambiguously to tested genes in both species. Across species-specific response categories, drying produced substantial shared transcript-abundance responses, whereas orthologs responding significantly to UVR in both species were comparatively sparse (Fig. 4A,B; Table S2).

**FIGURE 4.**
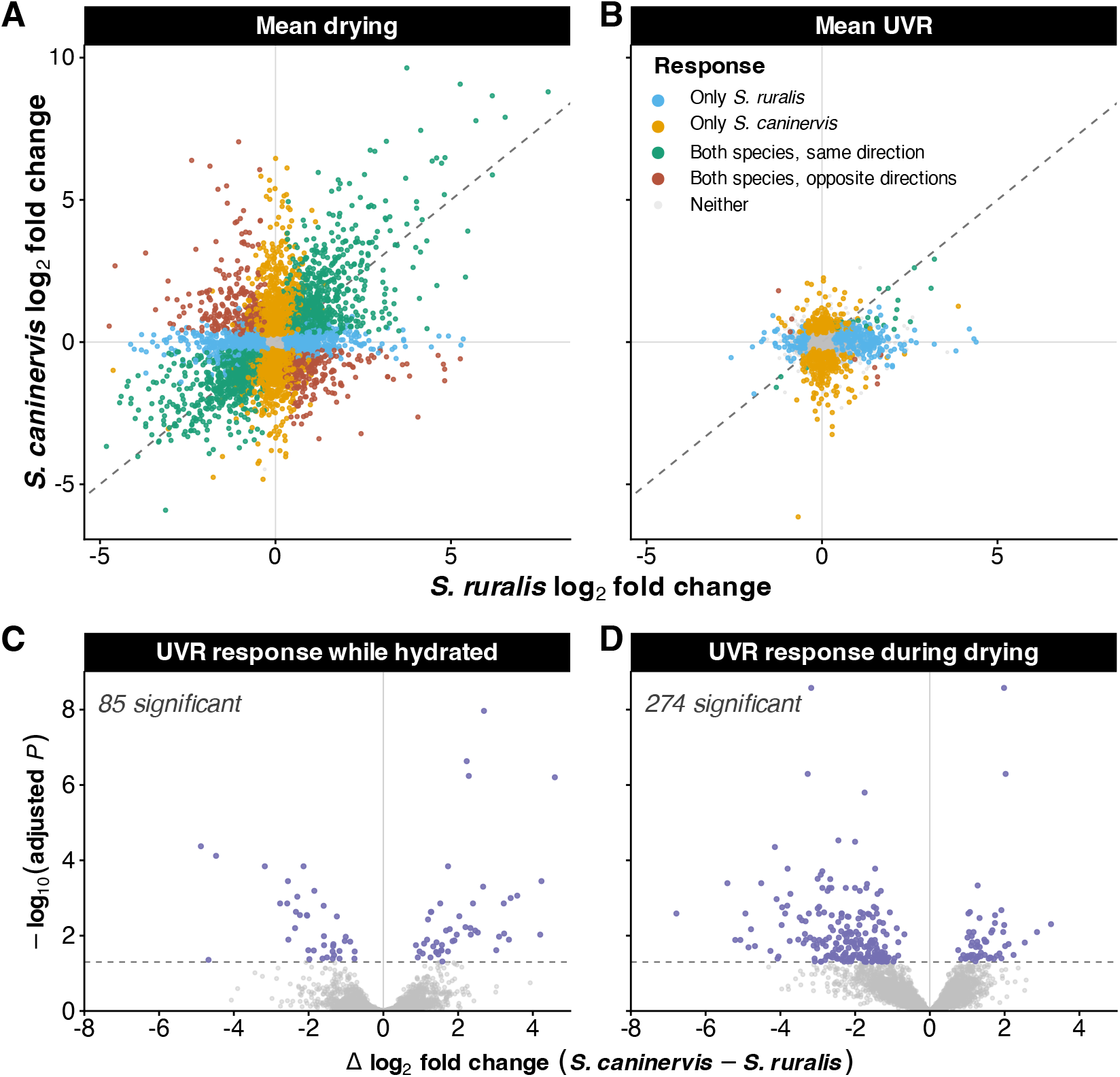
Drying responses are broadly shared across species, whereas UVR responses diverge more strongly during drying. (A) Cross-species comparison of mean drying log2 fold changes for mapped one-to-one orthologs. (B) Corresponding comparison of mean UVR log2 fold changes. In panels A and B, orthologs are colored by whether the response was significant in both species in the same or opposite direction, significant only in S. ruralis, significant only in S. caninervis, or not significant in either species; dashed lines show the 1:1 relationship. (C,D) Joint-model tests of the species difference in the UVR effect while hydrated and during drying, respectively. The joint DESeq2 model included 7,003 mapped orthogroups, of which 6,999 had testable adjusted P values for these contrasts, and identified 85 orthogroups with significant species differences in the UVR effect while hydrated and 274 during drying at Benjamini–Hochberg-adjusted P < 0.05. Positive values in panels C and D indicate a larger UVR effect in S. caninervis than in S. ruralis; the horizontal dashed line marks an adjusted P value of 0.05.

The joint orthogroup model directly tested whether the UVR response differed between species within each water-status condition (Fig. 4C,D). Among the 6,999 testable orthogroups, 85 showed a significant species difference in the UVR effect while hydrated, compared with 274 during drying. Species divergence in the UVR response was more extensive during drying than while hydrated.

### 5.4 Hydration and species strongly structure the soluble metabolome

The soluble and cell-wall-bound fractions differed in which environmental factors affected the most features. After filtering and quality control, the soluble datasets contained 4,261 features in negative mode and 6,392 in positive mode, while the cell-wall-bound datasets contained 1,789 and 4,972 features, respectively. One cell-wall-bound sample was excluded as a fraction-specific analytical outlier; the corresponding biological replicate behaved normally in the soluble datasets.

Across the four untargeted datasets, we analyzed 17,414 analytical features, including 10,653 soluble and 6,761 cell-wall-bound features. Direct GNPS library matching provided putative annotations for 572 features. Targeted m/z–retention-time matching added 99 features with a targeted putative compound annotation that lacked a direct GNPS annotation. In total, 654 features had an unconflicted putative compound assignment from at least one source. An additional 17 features had conflicting targeted and GNPS labels, and 22 had unresolved targeted candidate sets; these conflicts and ambiguities were retained and flagged in Table S9.

In the soluble fraction, hydration state produced the broadest response: 2,799 of 4,261 features in negative mode and 4,398 of 6,392 features in positive mode differed significantly between dry and hydrated tissues (Table S5). Species identity also affected a large proportion of the soluble metabolome, with 2,315 negative-mode and 3,779 positive-mode features significant for species. UVR treatment affected 1,009 negative-mode and 1,791 positive-mode features.

Species-dependent treatment responses were also widespread in the soluble fraction (Fig. 5A,B; Tables S5, S6). The species × hydration interaction was significant for 878 features in negative mode and 1,243 in positive mode, while the species × UVR interaction affected 706 and 1,355 features, respectively (Fig. 5B). Smaller but still detectable subsets of soluble features showed UVR × hydration interactions or three-way species × UVR × hydration interactions (Fig. 5B). Planned high-versus-low UVR contrasts showed that the breadth of the soluble UVR response varied with both species and water status (Fig. 5A; Table S6).

**FIGURE 5.**
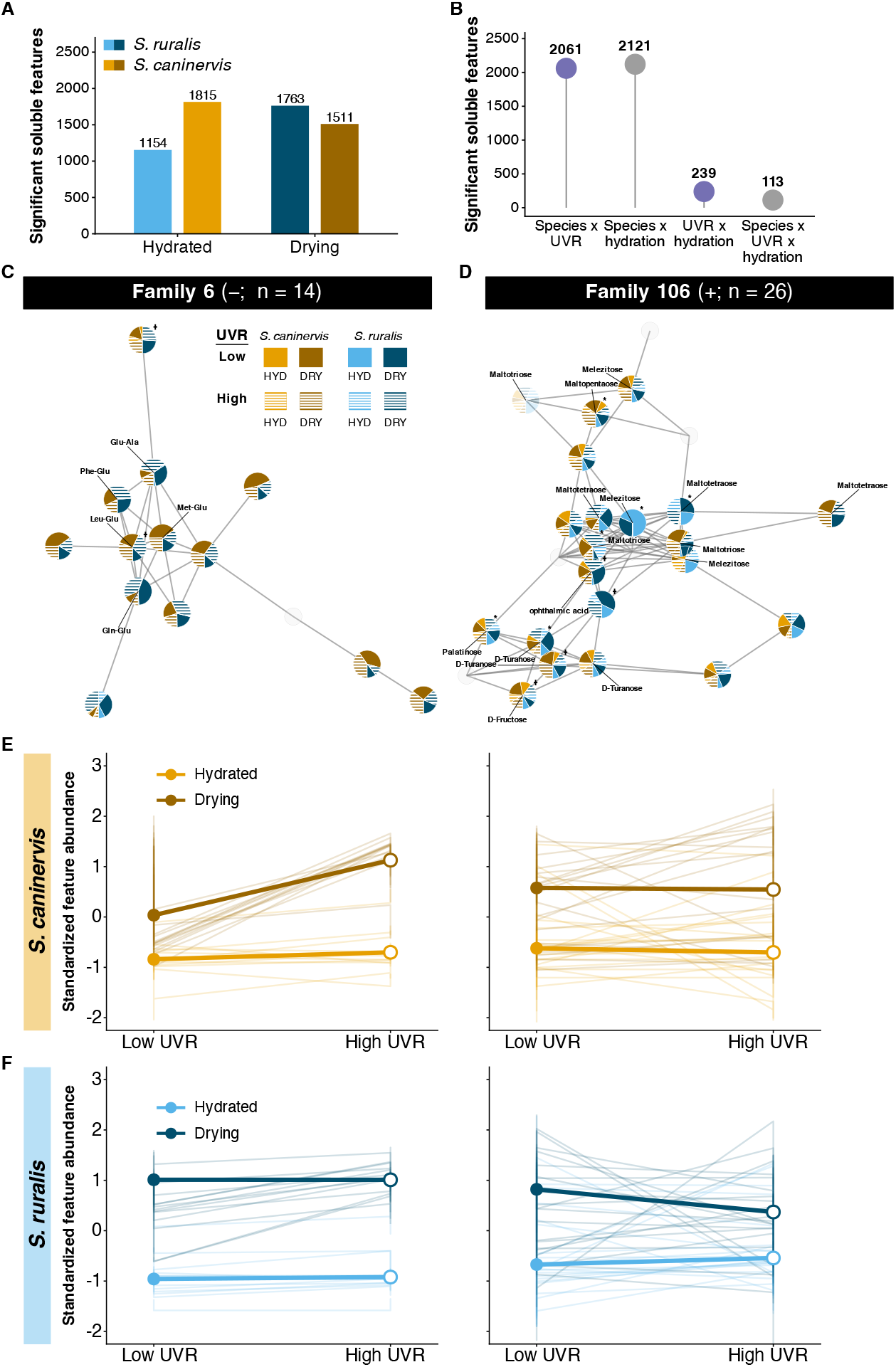
Soluble-metabolome UVR responses and representative molecular families. (A) Numbers of features differing significantly between high and low UVR within hydrated and drying tissue for each species, summed across separately analyzed ionization modes. (B) Numbers of features significant for each factorial interaction term. (C,D) GNPS molecular networks for negative-mode Family 6 (n = 14) and positive-mode Family 106 (n = 26); putative annotations are labeled where available. Opaque nodes were significant in at least one planned contrast or factorial-model term at Benjamini–Hochberg-adjusted P < 0.05. Asterisks indicate species × UVR interactions and daggers species × hydration interactions; symbols denote feature-level rather than family-level significance. (E,F) Standardized abundance profiles for Family 6 and Family 106 in S. caninervis and S. ruralis, respectively. Faint lines show individual features, and heavier lines with points show the median standardized abundance within each species × UVR × hydration group. Molecular-family membership is based on GNPS spectral similarity, and compound annotations are putative.

Several responsive features had putative metabolite annotations. Drying was associated with increased abundance of compounds including citrulline, tryptophan, ectoine, tyrosine, and ophthalmic acid, and decreased abundance of guanosine, lysine, putrescine, and several other annotated compounds. UVR-responsive annotated features included guanine, ectoine, and N-*α*-acetyl-lysine. A subset of annotated compounds also showed significant species × UVR or species × hydration interactions, including N-*α*-acetyl-lysine, nicotinic acid, isonicotinic acid, citrulline, and ectoine. These annotations are putative; annotation details are reported in Table S9, and feature-level effect patterns for the annotated soluble features are shown in Fig. S4.

### 5.5 Wall-associated chemistry shows a strong species-specific UVR signature

Wall-associated chemistry was proportionally more UVR responsive than the soluble fraction and showed a much stronger species × UVR interaction signature (Table S7). Species identity affected 791 of 1,789 negative-mode features and 2,569 of 4,972 positive-mode features. UVR treatment affected 434 negative-mode and 1,092 positive-mode features, whereas hydration affected comparatively fewer features: 194 and 636, respectively.

Nearly every significant interaction in the cell-wall-bound metabolome was species × UVR (Fig. 6B,C; Table S7). All 52 significant interaction features in negative mode were associated with species × UVR. In positive mode, 129 features were significant for species × UVR, whereas only one feature showed a UVR × hydration interaction and none showed species × hydration or three-way interactions. Across both ionization modes, 181 of 182 significant interaction features therefore represented species × UVR effects.

**FIGURE 6.**
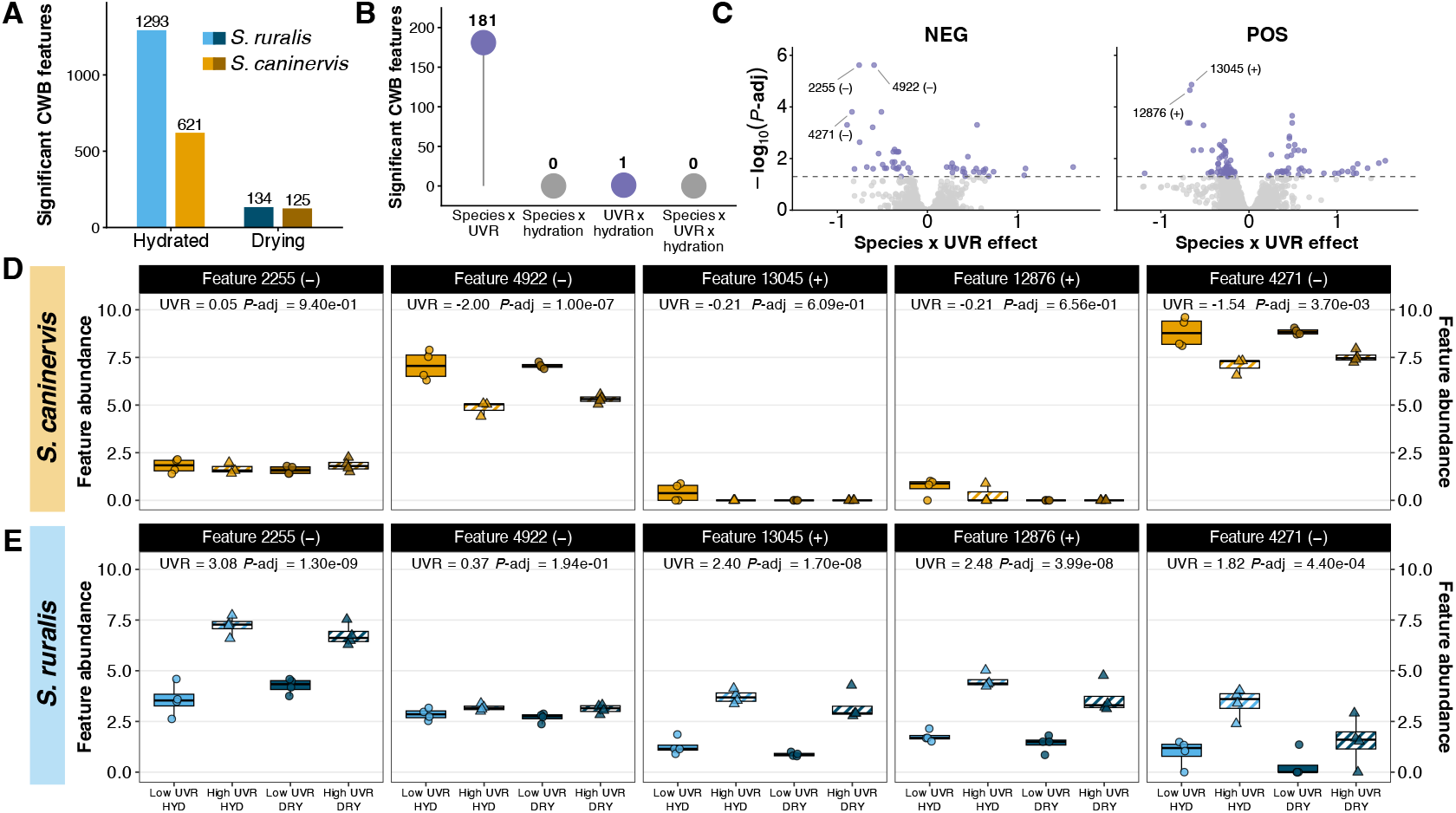
Cell-wall-bound UVR responses are concentrated in hydrated tissue and differ strongly between species. (A) Numbers of cell-wall-bound LC–MS features responding significantly to high versus low UVR within hydrated and drying tissue for each species; counts are summed across separately analyzed positive- and negative-ionization datasets for visual summary. (B) Numbers of features significant for each factorial interaction term across ionization modes; 181 of 182 significant interaction features were species × UVR effects, while one feature showed a UVR × hydration interaction and none showed species × hydration or three-way interactions. (C) Species × UVR interaction effect sizes and Benjamini–Hochberg-adjusted P values for negative- and positive-ionization features analyzed separately, with the five features displayed in panels D and E labeled. (D,E) Normalized feature abundance for those five species × UVR interaction features in S. caninervis and S. ruralis, respectively, across low and high UVR in hydrated (HYD) and drying (DRY) tissue. Symbols show biological replicates overlaid on boxplots; circles denote low UVR and triangles denote high UVR. Displayed UVR statistics in panels D and E are the species-specific UVR effect estimates and Benjamini–Hochberg-adjusted P values from the planned contrasts. Positive- and negative-ionization datasets were analyzed separately with limma, and significance was defined at Benjamini–Hochberg-adjusted P < 0.05.

Conditional UVR contrasts showed that wall-associated UVR responses were concentrated in hydrated tissue (Fig. 6A; Table S8). In negative mode, high versus low UVR affected 428 features in hydrated *S. ruralis* but only 59 during drying; corresponding values for *S. caninervis* were 121 and 55. Positive mode showed the same qualitative pattern, with 865 versus 75 significant features in hydrated and drying *S. ruralis*, and 500 versus 70 in *S. caninervis*. Selected species × UVR interaction features illustrate the underlying divergence: features 2255, 13045, and 12876 showed strong UVR responses in *S. ruralis* but little response in *S. caninervis*, feature 4922 showed the reverse pattern, and feature 4271 responded significantly in opposite directions in the two species (Fig. 6D,E).

GNPS molecular networks showed where species × UVR-responsive features occurred among spectrally related, largely unannotated features (Fig. 7). Positive-mode Family 121 contained 28 features, seven of which were significant for the species × UVR interaction (Fig. 7A). Negative-mode Family 44 showed a strongly species-skewed treatment distribution (Fig. 7B). Negative-mode Family 25 contained 28 features, including feature 2280, putatively annotated as (−)-catechin; two family members, features 4405 and 5992, were significant for the species × UVR interaction, whereas the catechin-annotated feature itself was not (Fig. 7C). Positive-mode Family 78 contained 19 features and one significant species × UVR feature (Fig. 7D). The treatment profiles of all four families differed between species, with family-level abundance patterns summarized separately for *S. caninervis* and *S. ruralis* (Fig. 7E,F).

**FIGURE 7.**
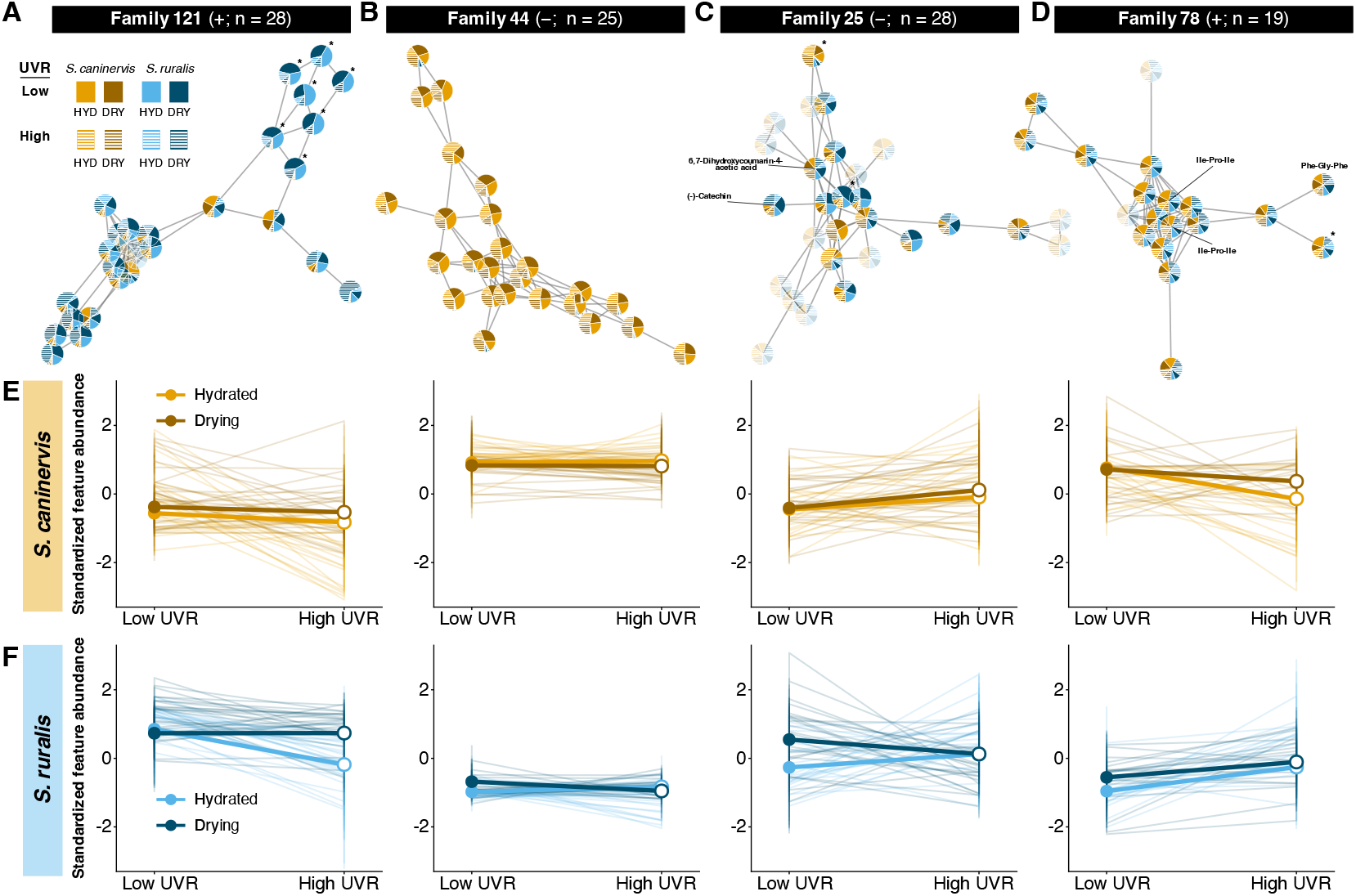
Species-specific UVR-responsive features occur within broader molecular families of wall-associated chemistry. (A–D) GNPS molecular families selected from the cell-wall-bound metabolome: positive-mode Family 121 (n = 28), negative-mode Family 44 (n = 25), negative-mode Family 25 (n = 28), and positive-mode Family 78 (n = 19), respectively. Node opacity distinguishes features significant in at least one planned contrast or factorial-model term at Benjamini–Hochberg-adjusted P < 0.05 from features not significant in any of those tests. Asterisks mark features significant specifically for the species × UVR interaction: seven in Family 121, none in Family 44, two in Family 25, and one in Family 78. These significance indicators are feature-level results rather than family-level significance tests. Putative annotations are shown where available, including (−)-catechin and 6,7-dihydroxycoumarin-4-acetic acid in Family 25 and peptide-like annotations in Family 78. (E,F) Standardized abundance profiles of all features in the four families for S. caninervis and S. ruralis, respectively, across low and high UVR in hydrated and drying tissue. Each feature was standardized across all biological samples in both species and treatments, so values indicate abundance relative to that feature’s overall mean and standard deviation rather than absolute abundance across features. Faint lines show individual features, and heavier lines with points show the median standardized abundance of family members within each species × UVR × hydration group. Molecular-family membership is based on GNPS spectral similarity, and compound annotations are putative.

## 6 DISCUSSION

Drying broadly reorganized the transcriptome and soluble metabolome in both species (Figs. 1D, 4A). This result builds on decades of work showing extensive molecular change across the drying–rehydration cycle in *Syntrichia*. Work in *S. ruralis* has long emphasized post-transcriptional regulation during desiccation, including changes in translational control and sequestration of mRNAs into messenger ribonucleoprotein particles, with substantial changes in protein synthesis becoming apparent upon rehydration (Oliver, 1991; Wood and Oliver, 1999). Within this broader regulatory response, we observed widespread changes in transcript abundance during drying (Fig. 4A), as has also been reported during dehydration in other desiccation-tolerant plants (Liu et al., 2018; Oliver et al., 2020).

The two species differed substantially in their soluble-metabolomic responses to drying (Fig. 5B), which may relate to previously reported differences in their desiccation physiology. Comparative work on the *Syntrichia ruralis* complex ranked *S. caninervis* as more desiccation tolerant than *S. ruralis*, with the difference attributed primarily to greater capacity to repair cellular damage after rehydration rather than to differences in damage during drying itself (Oliver et al., 1993). More recent work further shows that desiccation tolerance in *S. caninervis* is not uniformly constitutive: adult shoots are near constitutively tolerant, whereas protonemal tissue (filamentous juvenile moss tissue) requires much slower drying to achieve comparable recovery (Coe et al., 2021). These species-specific soluble-metabolomic responses therefore occur in a system where desiccation tolerance varies both among species and across life stages. Future work linking metabolomic responses during drying to subsequent recovery could test whether these molecular differences correspond to variation in desiccation tolerance.

The transcriptomic response to UVR depended on hydration state, and the pattern differed sharply between species. In *S. ruralis*, many more genes differed between UVR treatments during drying than while hydrated, whereas *S. caninervis* showed the opposite pattern (Fig. 3A). When we restricted the comparison to one-to-one orthologous genes shared between the two species, divergence in the UVR response was also greater during drying than while hydrated (Fig. 4C,D). These hydration-dependent responses also occurred in groups of genes with similar transcript-abundance patterns across UVR and hydration treatments within each species, rather than being limited to isolated genes (Fig. 3B,C).

Water availability sets the window in which these mosses can mount an active molecular response to UVR, and the consequences of that response may extend beyond that window into subsequent hydration cycles. In *S. ruralis*, repeated drying and rehydration increased tolerance to subsequent rapid drying, whereas prolonged continuous hydration reduced tolerance (Schonbeck and Bewley, 1981). Responses initiated while tissues are hydrated or during early drying may therefore persist through desiccation and influence recovery after rehydration. Drying can also change the radiation actually experienced by the plant: changes in leaf orientation and tissue structure alter how much incoming radiation reaches photosynthetic tissues (Seel et al., 1992). The hydration-dependent UVR responses observed here (Fig. 3) therefore occur as both the physiological capacity to respond and the physical exposure of tissues to radiation are changing.

One possibility is that UVR experienced during hydrated spring periods acts as a seasonal cue for the approaching summer dry season, inducing responses that persist into prolonged desiccation when plants remain exposed to intense UVR. In our previous acute-exposure study, ELIPs also showed a particularly strong UVR response in hydrated *S. caninervis*, including individually responsive ELIP transcripts (Ekwealor and Mishler, 2021). Here, the strongest annotated UVR × hydration interaction in *S. caninervis* was an ELIP transcript, whose abundance increased strongly under high UVR while hydrated but showed little additional UVR response during drying (Fig. 2A). The same transcript belonged to a 74-transcript module with a similar hydration-dependent response pattern (Module 3, Fig. 3B). In *Arabidopsis*, UV-B induction of ELIP1 and ELIP2 is UVR8 dependent, and circadian-clock components further regulate the UV-B response of ELIP1 (Hayami et al., 2015; Velanis et al., 2016; Takeuchi et al., 2014). The UVR8 photoreceptor and its basic mechanism are also conserved in the moss *Physcomitrium patens* (Soriano et al., 2018), making UVR8-mediated regulation a plausible, although untested, mechanism in *Syntrichia*.

UV-B also changes predictably with season: because UV-B increases more strongly with solar elevation than PAR, the UV-B:PAR ratio rises toward summer, making UV-B a potentially informative signal of seasonal progression (Robson et al., 2019). Concurrent temperature is a less reliable indicator of seasonal progression because it can vary independently of the longer-term seasonal trajectory (Tang et al., 2016). More broadly, UV-B can alter seasonal developmental processes in vascular plants, including bud phenology, and UVR8 signaling can interact directly with photoperiodic control of flowering (Strømme et al., 2015; Arongaus et al., 2018). Together with evidence that long-term UVR exclusion impairs recovery from desiccation in *S. caninervis* (Ekwealor et al., 2021), these results strengthen the hypothesis that UVR experienced during hydrated periods may help prepare plants for the prolonged desiccation that follows. A direct test could manipulate UVR during the hydrated period and then measure ELIP responses, subsequent desiccation tolerance, and photosynthetic recovery after rehydration.

The *S. ruralis* response may instead reflect greater reliance on responses mounted during drying itself. We expected its UVR response to be strongest while tissues were hydrated, yet substantially more genes differed between UVR treatments during drying. Because *S. ruralis* occupies generally more mesic habitats than *S. caninervis*, one possibility is that it more often experiences drying slowly enough to maintain physiological activity while water content declines, allowing UVR-associated responses to develop during the drying process rather than being established in advance. By contrast, plants exposed to rapid and prolonged drying may benefit more from responses induced while tissues are still fully hydrated.

UVR and hydration reshaped the soluble metabolome differently in the two species (Fig. 5A,B). The annotated soluble features included compounds associated with amino-acid, osmolyte, and redox-related chemistry, consistent with shifts in soluble metabolism during UVR exposure and drying. Within the highlighted molecular families, spectrally related features often showed similar treatment-associated abundance patterns even when most remained unannotated, indicating that these responses extended across broader groups of related chemistry than the small subset of compounds that could be identified (Fig. 5C–F).

UVR effects on wall-associated chemistry diverged sharply between the two species (Fig. 6). Nearly all significant interactions reflected species × UVR (181 of 182), whereas interactions involving hydration were rare, indicating that the two species differed primarily in how wall-associated chemistry responded to UVR rather than in how hydration modified that response. Because wall-associated compounds can remain in place as tissues dry, these species-specific responses could contribute to chemical differences that persist beyond the hydrated period, potentially offering passive UVR protection.

Persistent wall-associated photoprotection may come with physiological costs, however. UVR-screening compounds at or near the tissue surface are positioned to intercept incoming radiation and can remain in place as tissues lose water (Clarke and Robinson, 2008), and wall-associated phenolics can constitute a substantial fraction of UVR-screening chemistry in bryophytes (Taipale and Huttunen, 2002; Clarke and Robinson, 2008; Robinson and Waterman, 2014; Waterman et al., 2017). Across bryophytes, greater concentrations of wall-bound UVR-absorbing compounds are associated with lower maximum photosynthetic capacity, consistent with a potential cost through reduced CO_2_ diffusion across phenolic-rich walls (Perera Castro et al., 2026; Ekwealor and Simonin, 2026). If maintaining persistent wall-associated protection carries such a cost, species that commonly remain physiologically active during gradual drying may benefit from accumulating protective compounds as water is lost rather than maintaining them in advance. This possibility could help explain why *S. ruralis*, which occupies generally more mesic habitats, showed its broadest UVR-associated transcriptomic response during drying, whereas *S. caninervis* showed a stronger response while hydrated.

One molecular family provides a tentative chemical clue to the wall-associated UVR response. The negative-mode family highlighted in Figure 7C contained two species × UVR-responsive features and a feature putatively annotated as (−)-catechin, although the catechin-annotated feature itself was not significant for the interaction (Table S9). Catechin is a flavan-3-ol flavonoid that has also been detected in other mosses (Naznin et al., 2025), while flavonoid and other phenolic chemistry is well established in bryophyte UV responses (Soriano et al., 2019). Although neighboring nodes cannot be assigned as catechin derivatives from spectral similarity alone, their clustering raises the possibility that this species-specific UVR response involves related flavonoid chemistry.

The ecological contrast between the species provides useful context for these differences but did not fully predict them. *Syntrichia caninervis* is more strongly associated with arid habitats and has greater reported desiccation tolerance than *S. ruralis*, which occupies a broader range of moisture regimes (Mishler, 2007; Oliver et al., 1993). Our results extend this divergence in water relations by showing that the species also differ in when UVR-associated responses occur and in which molecular compartments they are strongest. Drying broadly reorganized the transcriptome and soluble metabolome, transcriptomic UVR responsiveness shifted in opposite directions during drying in the two species, and wall-associated chemistry showed strong species-specific UVR responses that were much broader in hydrated than drying tissue. Water availability therefore shapes not only the magnitude of molecular responses to UVR, but when and where those responses occur, raising the question of whether changes established during hydrated periods persist through desiccation and influence later performance.

## Acknowledgements

The authors would like to thank Carl J. Rothfels, Cindy V. Looy, and Mary K. Firestone for their feedback on earlier versions of this manuscript. The authors acknowledge the technical skills of Kate Guill and Anderson Silva in the preparation of RNA-Seq libraries. We also thank Cindy V. Looy, Ivo A. P. Duijnstee, and Jeffrey P. Benca for their assistance with measuring UV-B flux from our experimental lamps.

## Author Contributions

JTBE, BDM, MJO, and TN designed the research. JTBE, ATS, and SK performed the research. JTBE and SK performed data collection. JTBE and SK performed data analyses. All authors contributed to interpretation and writing.

## Data Availability

Code and processed data supporting the analyses are available at https://github.com/jenna-tb-ekwealor/UVR_DRY_multi-omics. Raw transcriptomic sequence data will be deposited in the NCBI Sequence Read Archive, and additional metabolomic and transcriptomic data will be deposited in a public data repository before publication. Accession numbers and persistent identifiers will be added upon deposition.

## Supporting Information

**FIGURE S1.**
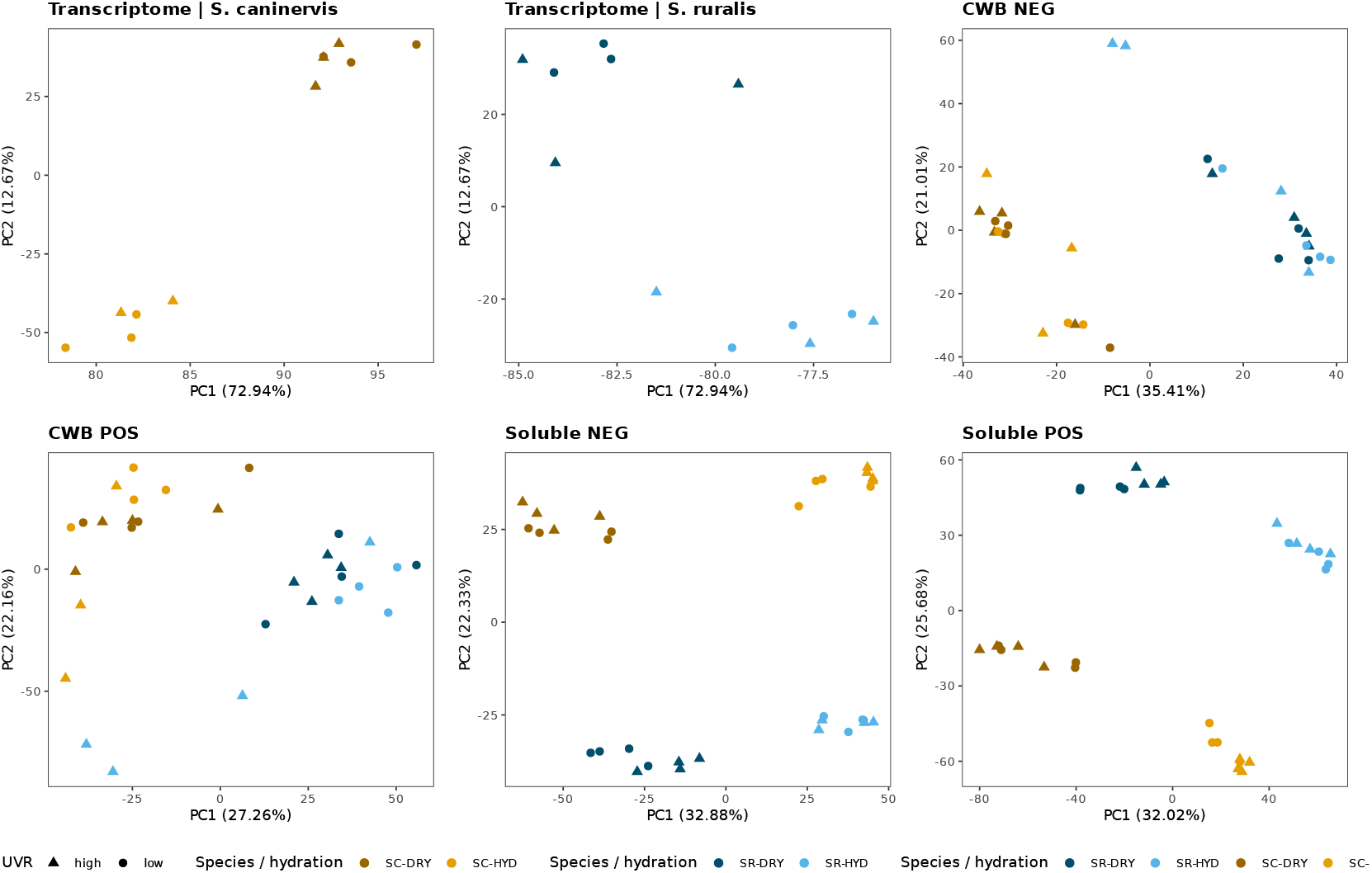
Separate principal component analyses for transcriptomic and metabolomic datasets. The top-left and top-middle panels show variance-stabilized transcript abundance for S. caninervis and S. ruralis, respectively. The remaining panels show cell-wall-bound (CWB) and soluble metabolomic datasets analyzed separately in negative- and positive-ionization modes. Point color indicates species and hydration state, and point shape indicates UVR treatment. Axis labels report the percentage of variance explained by each principal component.

**FIGURE S2.**
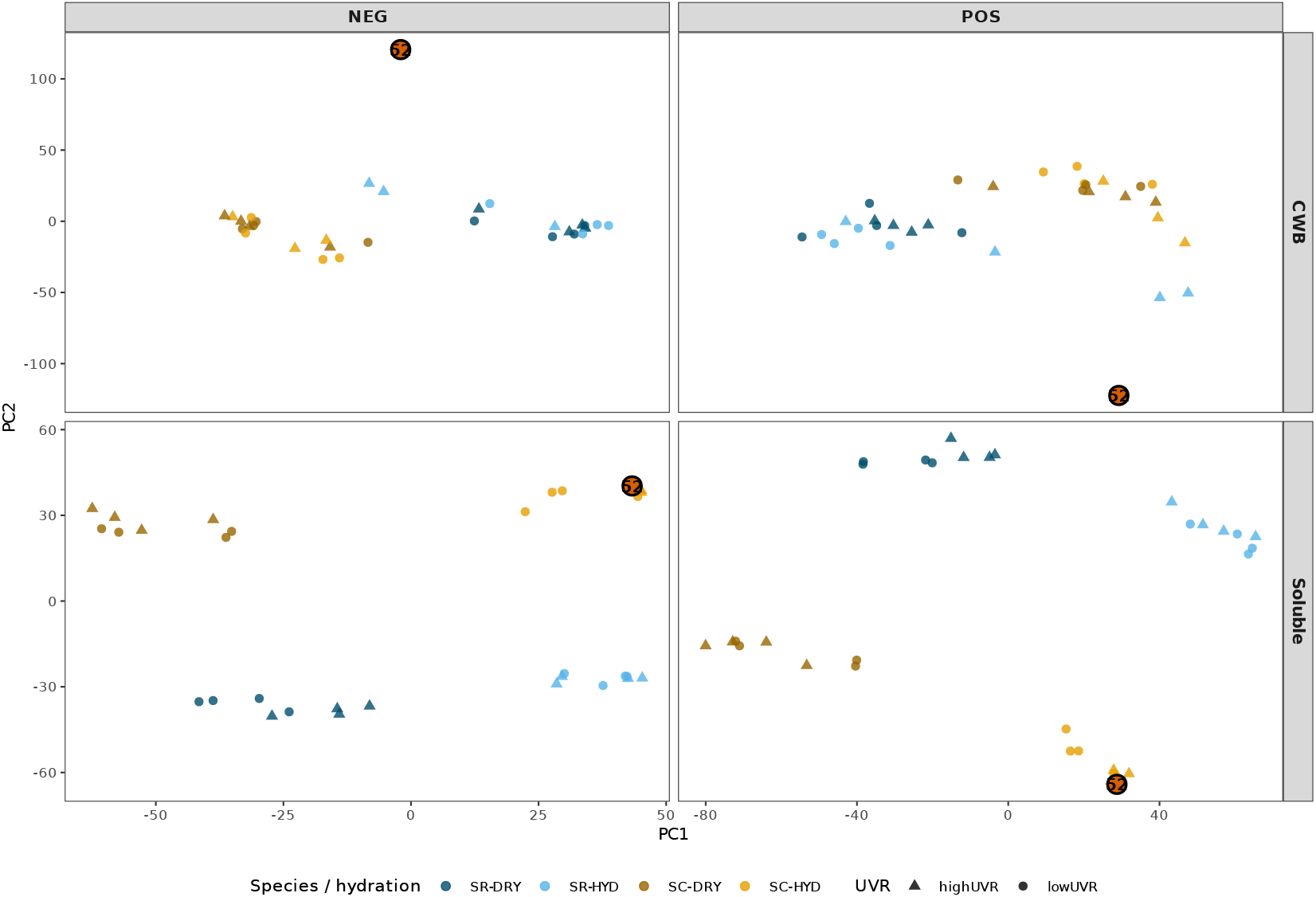
Diagnostic PCA views for metabolomics sample 52. Sample 52 is highlighted in red in negative- and positive-ionization PCA space for the cell-wall-bound and soluble fractions. The sample was a strong multivariate outlier in both cell-wall-bound datasets and was excluded from primary cell-wall-bound analyses, whereas the corresponding biological replicate was retained in the soluble datasets. Colors indicate species and hydration state, and symbols indicate UVR treatment.

**FIGURE S3.**
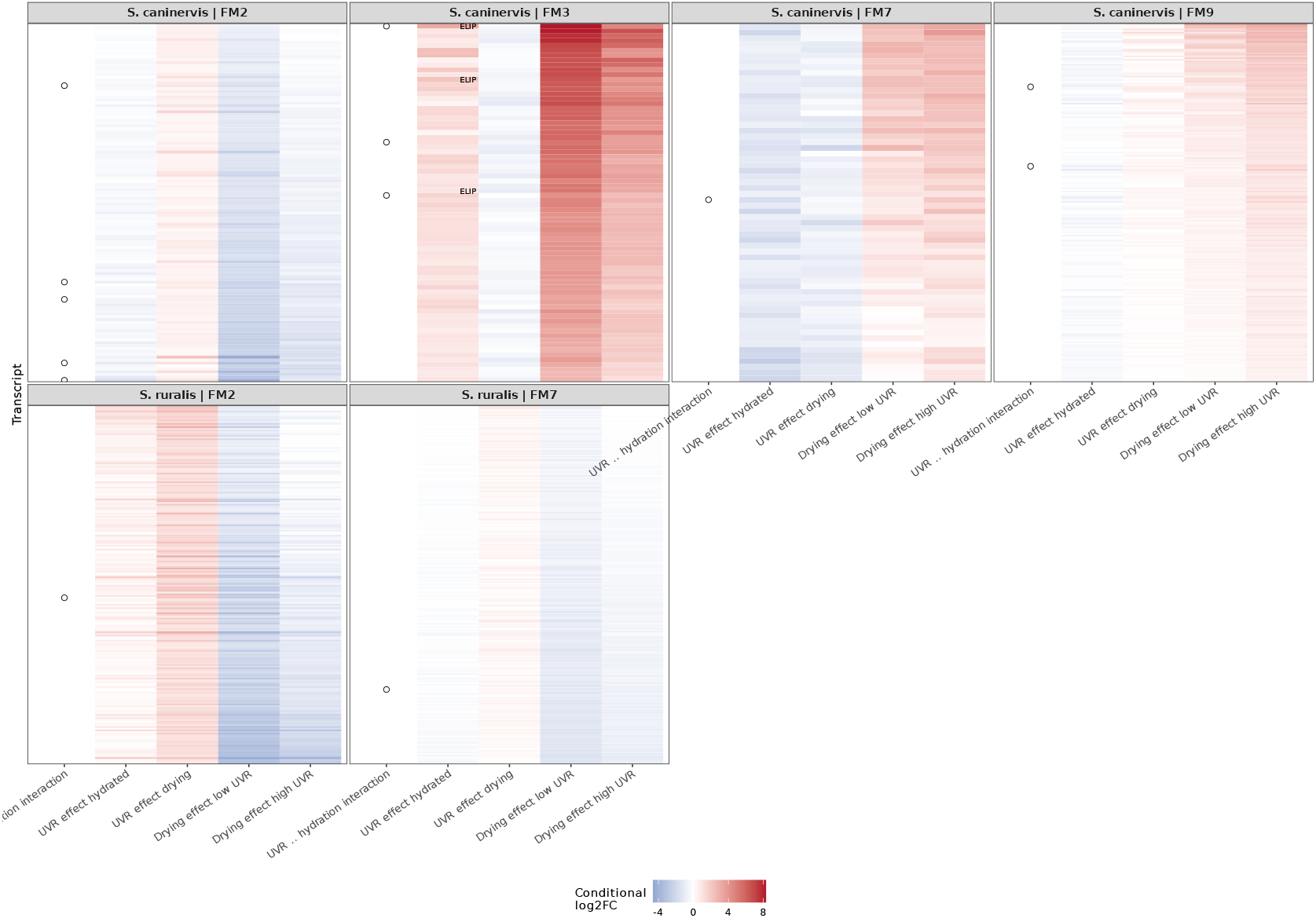
Conditional response structure of retained transcript-response modules. Rows are transcripts assigned to the retained high-coherence modules in S. caninervis and S. ruralis, and heatmap color shows the modeled log2 fold change for the UVR effect while hydrated, UVR effect during drying, drying effect under low UVR, and drying effect under high UVR. The left-hand marker denotes transcripts significant for the UVR × hydration interaction, whereas black ticks within the heatmap mark significance for the matching conditional contrast. Three distinct ELIP-annotated transcripts are labeled within S. caninervis Module 3. Module membership, row order, and effect estimates are the same as in the primary module analysis.

**FIGURE S4.**
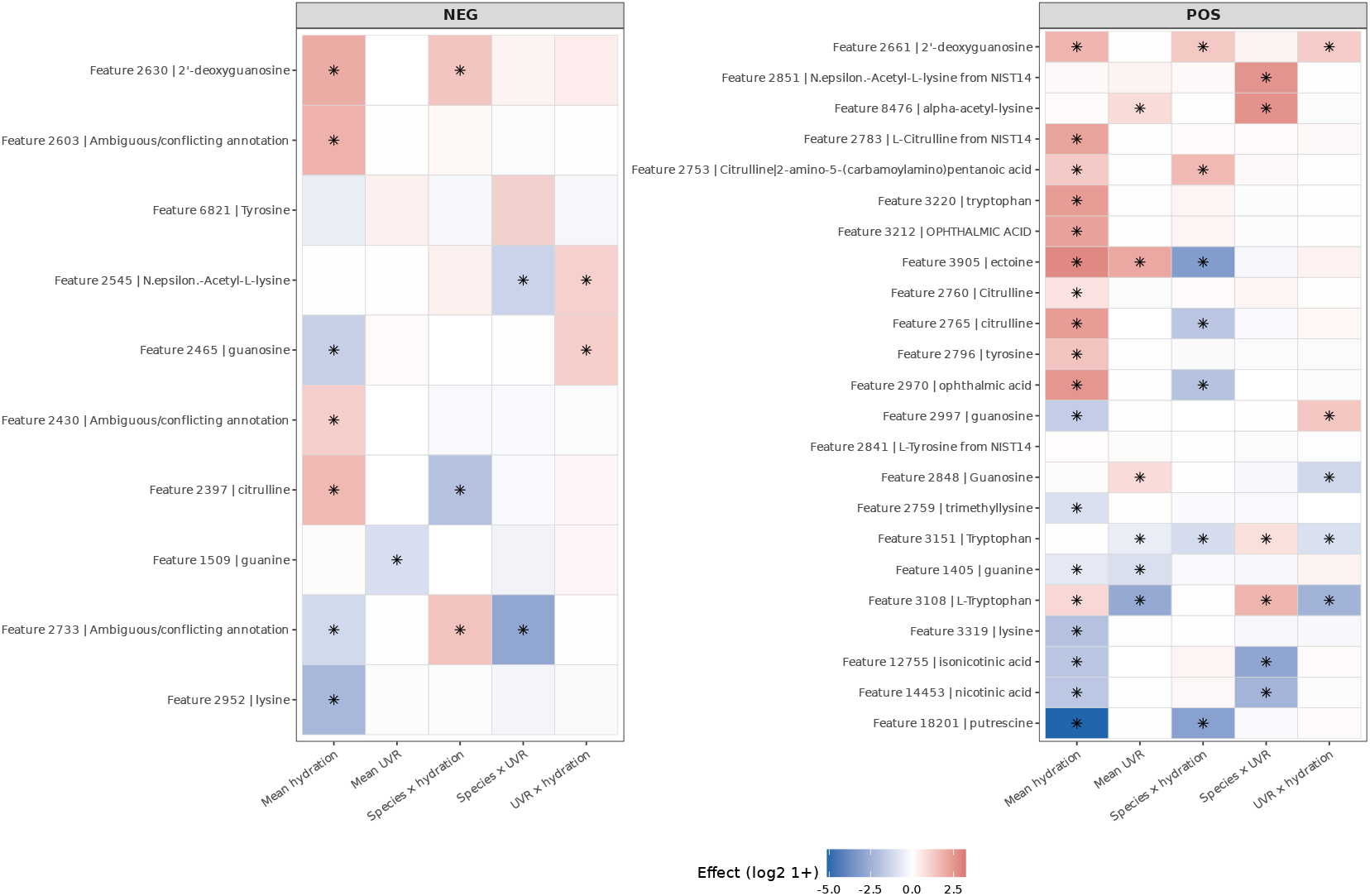
Treatment responses of soluble-metabolomic features with annotation evidence. Negative- and positive-ionization features are shown separately, with feature IDs retained because the same putative annotation can occur for multiple distinct LC–MS features. Cells show modeled effect sizes for mean hydration, mean UVR, species × hydration, species × UVR, and UVR × hydration effects. Asterisks mark effects significant at Benjamini–Hochberg-adjusted P < 0.05. Assigned compound identities are putative; ambiguous or conflicting annotations are shown without a compound identity.

**FIGURE S5.**
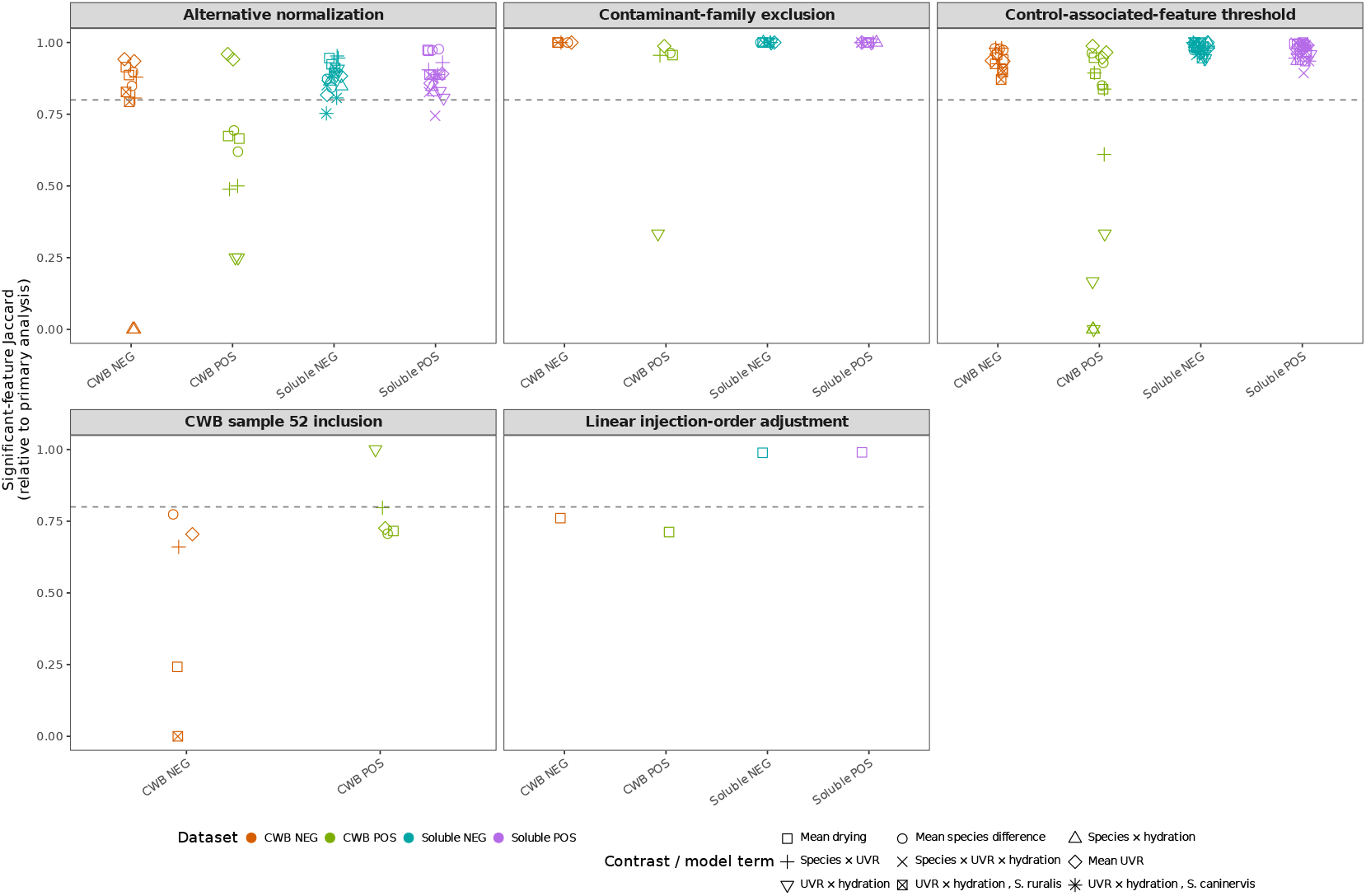
Robustness and sensitivity of metabolomic differential-abundance results. Points show Jaccard similarity in the set of significant features between the primary analysis and sensitivity analyses using alternative normalization, conservative contaminant-family exclusion, an alternative control-associated-feature threshold, inclusion of cell-wall-bound sample 52, and linear adjustment for injection order. Colors identify the four analytical datasets, and point shapes identify the corresponding contrast or factorial-model term. Only finite Jaccard values are plotted. The horizontal dashed line at 0.8 is a visual reference only and is not an inferential or decision threshold.

**TABLE S1.**
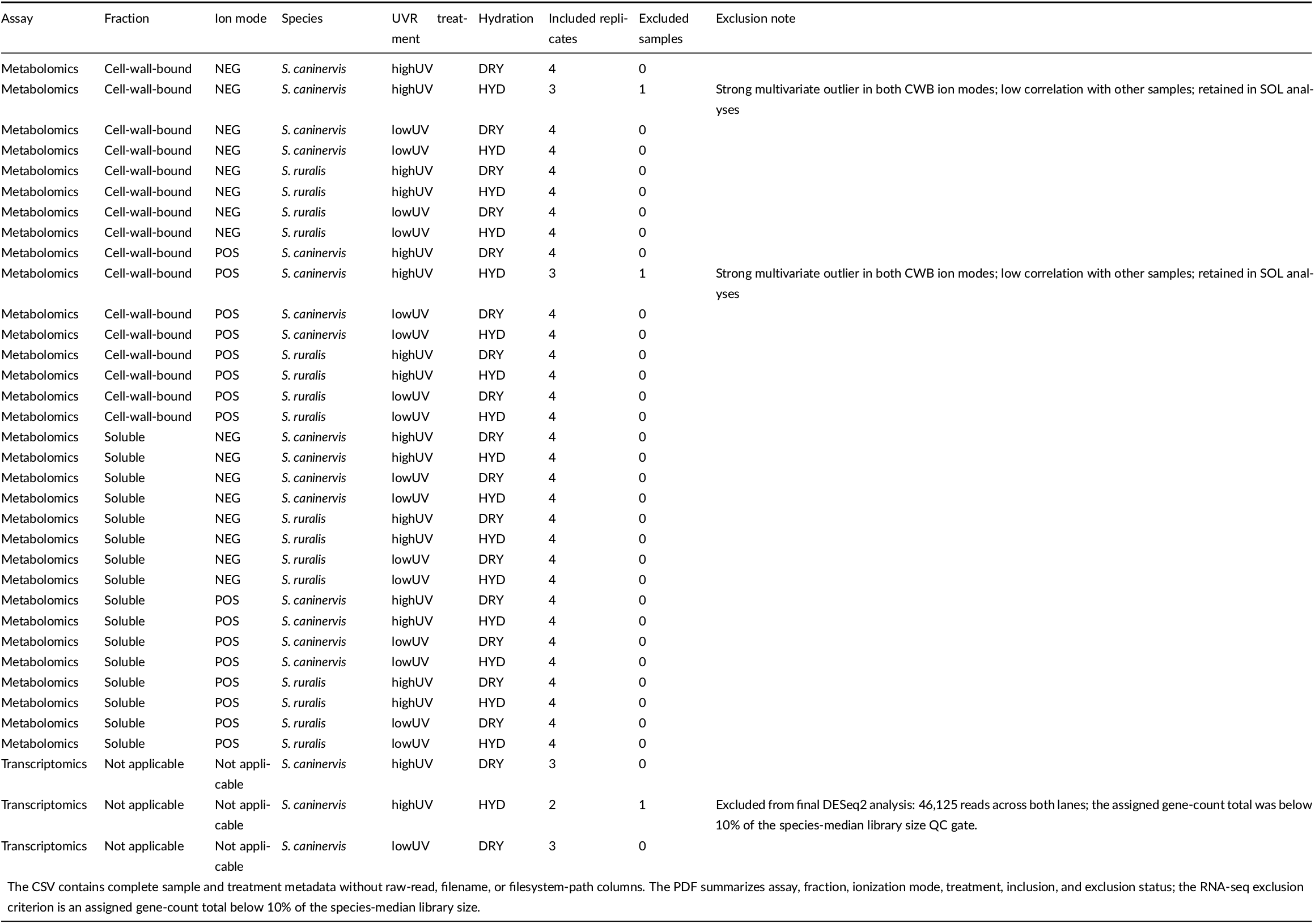
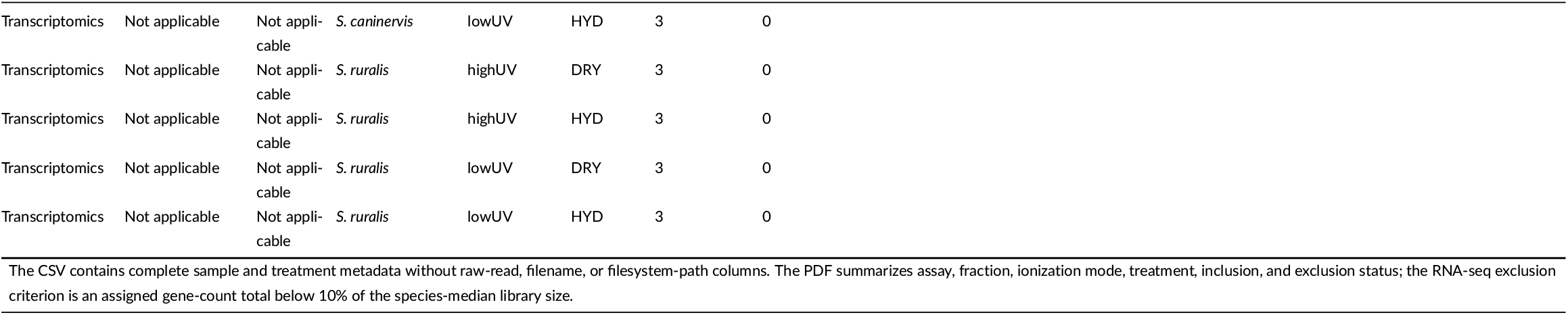
Sample design and primary-analysis inclusion summary.

**TABLE S2.**
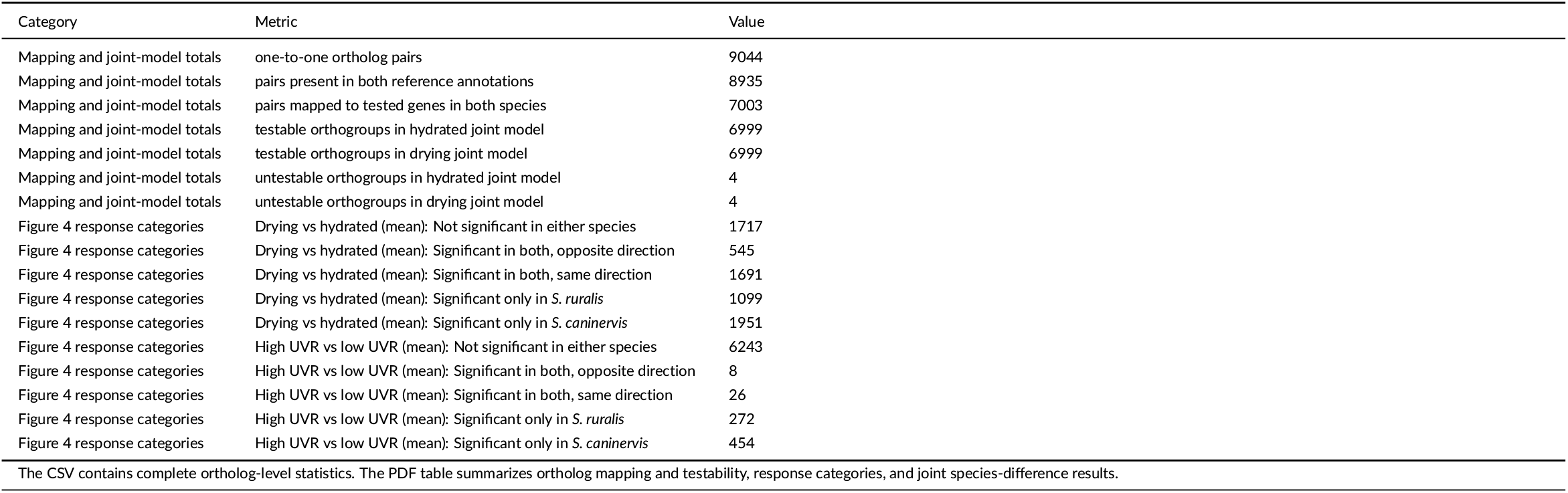
Cross-species ortholog analysis summary.

| Category | Metric | Value |
| --- | --- | --- |
| Mapping and joint-model totals | one-to-one ortholog pairs | 9044 |
| Mapping and joint-model totals | pairs present in both reference annotations | 8935 |
| Mapping and joint-model totals | pairs mapped to tested genes in both species | 7003 |
| Mapping and joint-model totals | testable orthogroups in hydrated joint model | 6999 |
| Mapping and joint-model totals | testable orthogroups in drying joint model | 6999 |
| Mapping and joint-model totals | untestable orthogroups in hydrated joint model | 4 |
| Mapping and joint-model totals | untestable orthogroups in drying joint model | 4 |
| Figure 4 response categories | Drying vs hydrated (mean): Not significant in either species | 1717 |
| Figure 4 response categories | Drying vs hydrated (mean): Significant in both, opposite direction | 545 |
| Figure 4 response categories | Drying vs hydrated (mean): Significant in both, same direction | 1691 |
| Figure 4 response categories | Drying vs hydrated (mean): Significant only in *S. ruralis* | 1099 |
| Figure 4 response categories | Drying vs hydrated (mean): Significant only in *S. caninervis* | 1951 |
| Figure 4 response categories | High UVR vs low UVR (mean): Not significant in either species | 6243 |
| Figure 4 response categories | High UVR vs low UVR (mean): Significant in both, opposite direction | 8 |
| Figure 4 response categories | High UVR vs low UVR (mean): Significant in both, same direction | 26 |
| Figure 4 response categories | High UVR vs low UVR (mean): Significant only in *S. ruralis* | 272 |
| Figure 4 response categories | High UVR vs low UVR (mean): Significant only in *S. caninervis* | 454 |

The CSV contains complete ortholog-level statistics. The PDF table summarizes ortholog mapping and testability, response categories, and joint species-difference results.

**TABLE S3.**
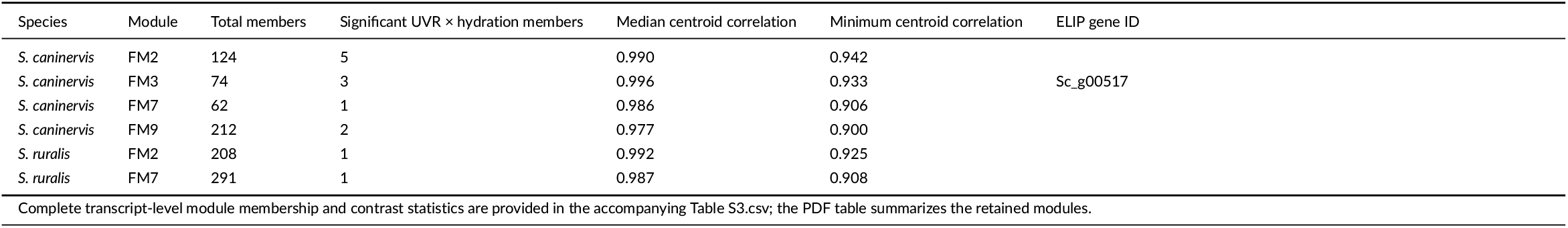
Retained transcript-response module summary.

**TABLE S4.** Transcript-response module summary.

**Significant members by response test**
| Species | Module | UVR × hydration in-<br>teraction | UVR effect, hydrated | UVR effect, drying | Drying, low UVR | Drying, high UVR |
| --- | --- | --- | --- | --- | --- | --- |
| *S. caninervis* | FM2 | 5/124 (4.0%) | 2/124 (1.6%) | 6/124 (4.8%) | 120/124 (96.8%) | 46/124 (37.1%) |
| *S. caninervis* | FM3 | 3/74 (4.1%) | 8/74 (10.8%) | 0/74 (0.0%) | 74/74 (100.0%) | 67/74 (90.5%) |
| *S. caninervis* | FM7 | 1/62 (1.6%) | 24/62 (38.7%) | 17/62 (27.4%) | 38/62 (61.3%) | 44/62 (71.0%) |
| *S. caninervis* | FM9 | 2/212 (0.9%) | 3/212 (1.4%) | 3/212 (1.4%) | 101/212 (47.6%) | 180/212 (84.9%) |
| *S. ruralis* | FM2 | 1/208 (0.5%) | 1/208 (0.5%) | 94/208 (45.2%) | 174/208 (83.7%) | 66/208 (31.7%) |
| *S. ruralis* | FM7 | 1/291 (0.3%) | 0/291 (0.0%) | 3/291 (1.0%) | 189/291 (64.9%) | 42/291 (14.4%) |

Significance was assessed at BH-adjusted $P < 0.05$ within the corresponding response test.

**TABLE S5.**
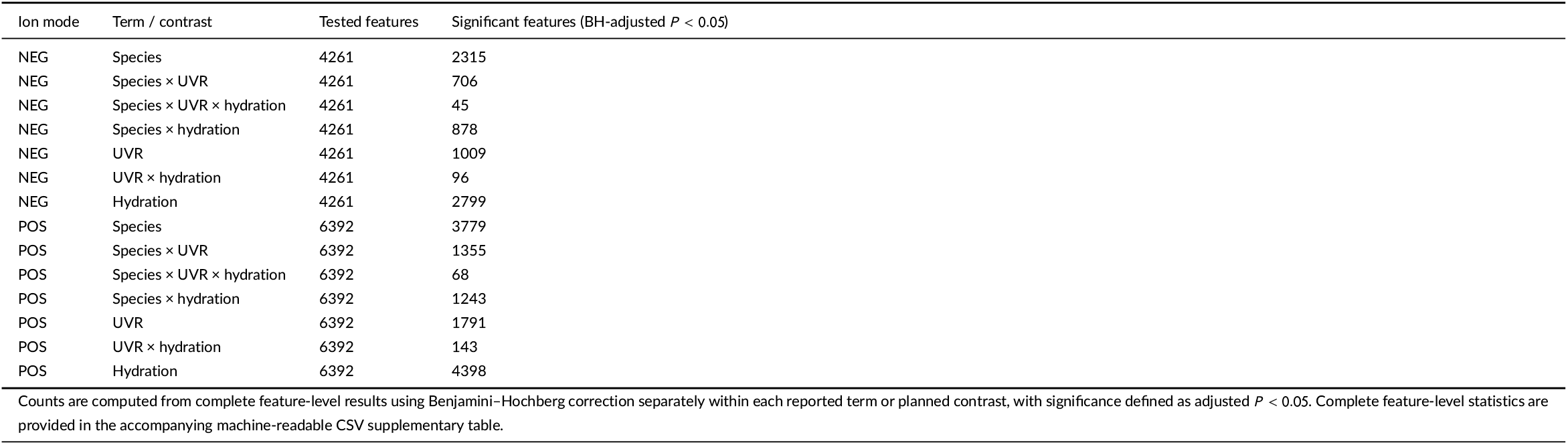
Soluble metabolome factorial-term result.

| Ion mode | Term / contrast | Tested features | Significant features (BH-adjusted $P < 0.05$) |
| --- | --- | --- | --- |
| NEG | Species | 4261 | 2315 |
| NEG | Species × UVR | 4261 | 706 |
| NEG | Species × UVR × hydration | 4261 | 45 |
| NEG | Species × hydration | 4261 | 878 |
| NEG | UVR | 4261 | 1009 |
| NEG | UVR × hydration | 4261 | 96 |
| NEG | Hydration | 4261 | 2799 |
| POS | Species | 6392 | 3779 |
| POS | Species × UVR | 6392 | 1355 |
| POS | Species × UVR × hydration | 6392 | 68 |
| POS | Species × hydration | 6392 | 1243 |
| POS | UVR | 6392 | 1791 |
| POS | UVR × hydration | 6392 | 143 |
| POS | Hydration | 6392 | 4398 |

Counts are computed from complete feature-level results using Benjamini–Hochberg correction separately within each reported term or planned contrast, with significance defined as adjusted $P < 0.05$ . Complete feature-level statistics are provided in the accompanying machine-readable CSV supplementary table.

**TABLE S6.** Soluble metabolome planned-contrast results.

| Ion mode | Term / contrast | Tested features | Significant features (BH-adjusted $P < 0.05$) |
| --- | --- | --- | --- |
| NEG | Drying vs hydrated: *S. caninervis* high | 4261 | 2128 |
| NEG | Drying vs hydrated: *S. caninervis* low | 4261 | 1629 |
| NEG | Drying vs hydrated: *S. ruralis* high | 4261 | 1100 |
| NEG | Drying vs hydrated: *S. ruralis* low | 4261 | 1781 |
| NEG | Drying vs hydrated: in *S. caninervis* | 4261 | 2427 |
| NEG | Drying vs hydrated: in *S. ruralis* | 4261 | 2147 |
| NEG | Drying vs hydrated: mean | 4261 | 2799 |
| NEG | High UVR vs low UVR: *S. caninervis* (drying) | 4261 | 509 |
| NEG | High UVR vs low UVR: *S. caninervis* (hydrated) | 4261 | 795 |
| NEG | High UVR vs low UVR: *S. ruralis* (drying) | 4261 | 510 |
| NEG | High UVR vs low UVR: *S. ruralis* (hydrated) | 4261 | 381 |
| NEG | High UVR vs low UVR: in *S. caninervis* | 4261 | 973 |
| NEG | High UVR vs low UVR: in *S. ruralis* | 4261 | 699 |
| NEG | Species *S. caninervis* vs *S. ruralis* mean | 4261 | 2315 |
| NEG | Species × Hydration | 4261 | 878 |
| NEG | Species × UVR | 4261 | 706 |
| NEG | Species × UVR × Hydration | 4261 | 45 |
| NEG | UVR High vs Low mean | 4261 | 1009 |
| NEG | UVR × Hydration | 4261 | 96 |
| NEG | UVR × Hydration in *S. caninervis* | 4261 | 88 |
| NEG | UVR × Hydration in *S. ruralis* | 4261 | 53 |
| POS | Drying vs hydrated: *S. caninervis* high | 6392 | 3329 |
| POS | Drying vs hydrated: *S. caninervis* low | 6392 | 2488 |
| POS | Drying vs hydrated: *S. ruralis* high | 6392 | 2068 |
| POS | Drying vs hydrated: *S. ruralis* low | 6392 | 3126 |
| POS | Drying vs hydrated: in *S. caninervis* | 6392 | 3831 |
| POS | Drying vs hydrated: in *S. ruralis* | 6392 | 3538 |
| POS | Drying vs hydrated: mean | 6392 | 4398 |
| POS | High UVR vs low UVR: *S. caninervis* (drying) | 6392 | 1002 |
| POS | High UVR vs low UVR: *S. caninervis* (hydrated) | 6392 | 1020 |
| POS | High UVR vs low UVR: *S. ruralis* (drying) | 6392 | 1253 |
| POS | High UVR vs low UVR: *S. ruralis* (hydrated) | 6392 | 773 |
| POS | High UVR vs low UVR: in *S. caninervis* | 6392 | 1544 |
| POS | High UVR vs low UVR: in *S. ruralis* | 6392 | 1599 |
| POS | Species *S. caninervis* vs *S. ruralis* mean | 6392 | 3779 |
| POS | Species × Hydration | 6392 | 1243 |
| POS | Species × UVR | 6392 | 1355 |
| POS | Species × UVR × Hydration | 6392 | 68 |
| POS | UVR High vs Low mean | 6392 | 1791 |
| POS | UVR × Hydration | 6392 | 143 |
| POS | UVR × Hydration in *S. caninervis* | 6392 | 222 |
| POS | UVR × Hydration in *S. ruralis* | 6392 | 90 |

Counts are computed from complete feature-level results using Benjamini–Hochberg correction separately within each reported term or planned contrast, with significance defined as adjusted $P < 0.05$ . Complete feature-level statistics are provided in the accompanying machine-readable CSV supplementary table.

**TABLE S7.**
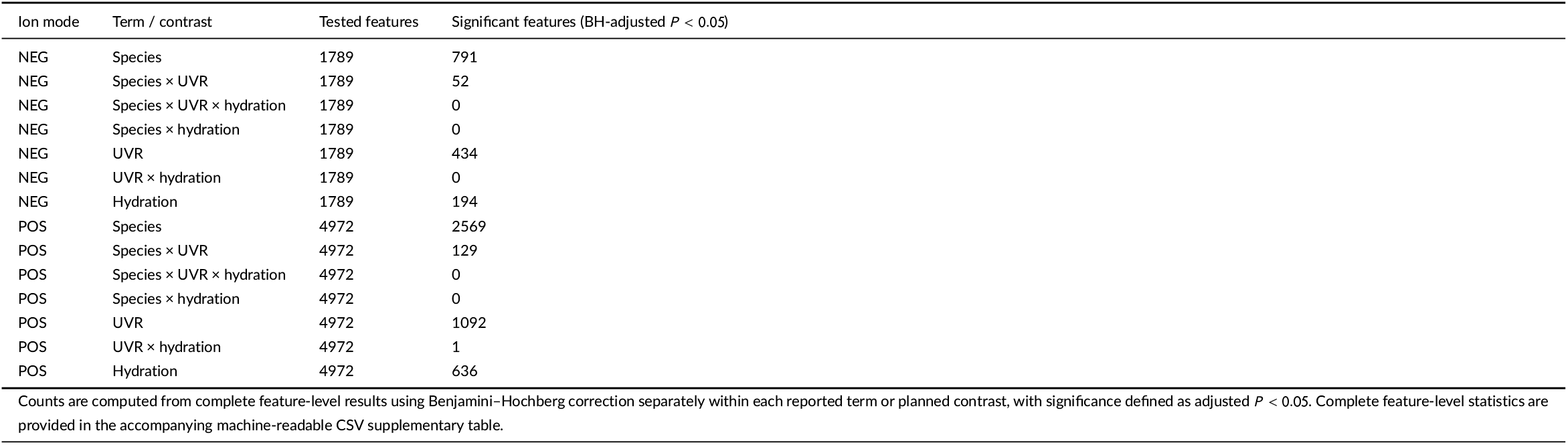
Cell-wall-bound metabolome factorial-term results.

**TABLE S8.**
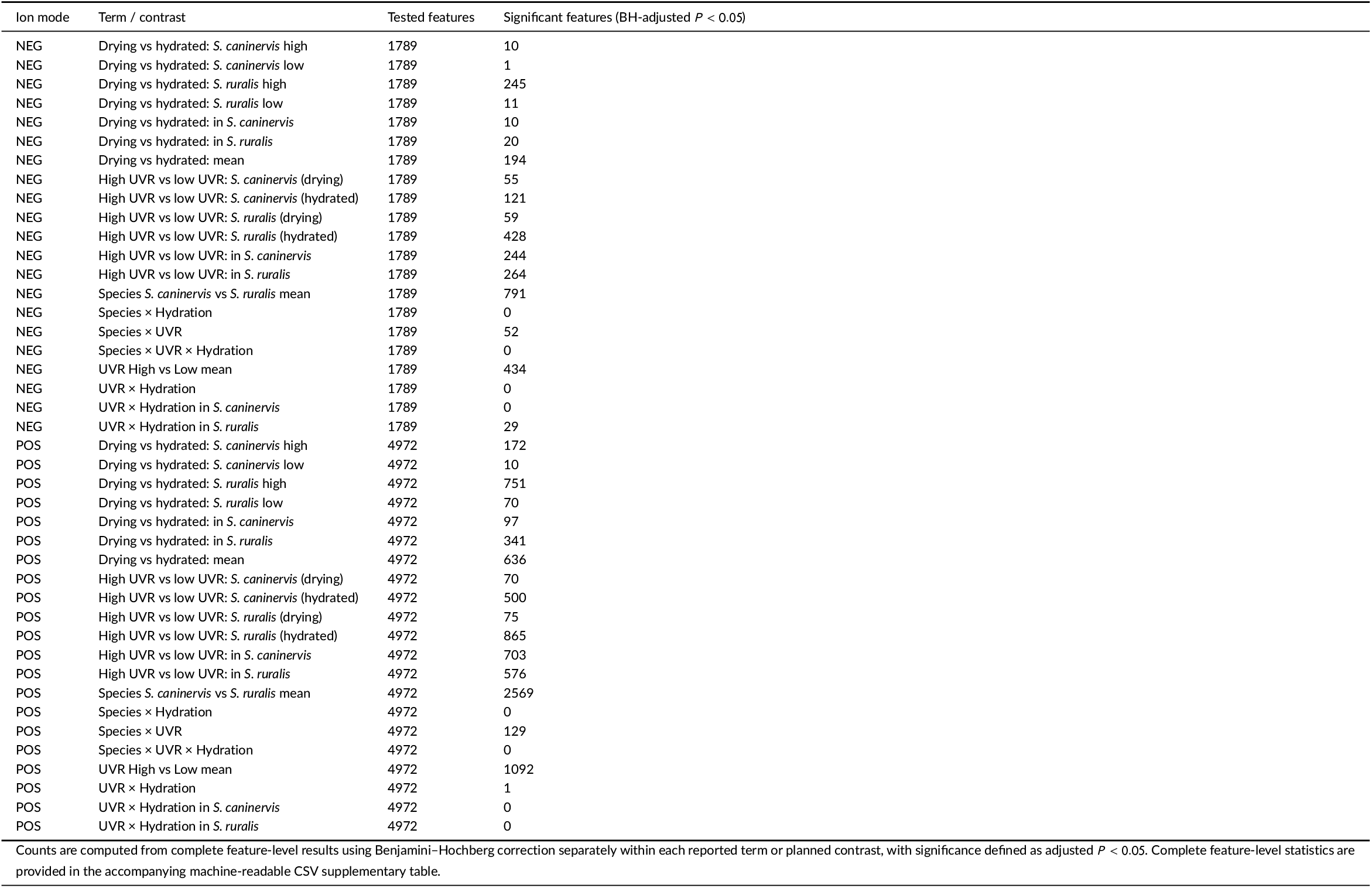
Cell-wall-bound metabolome planned-contrast results.

**TABLE S9.**
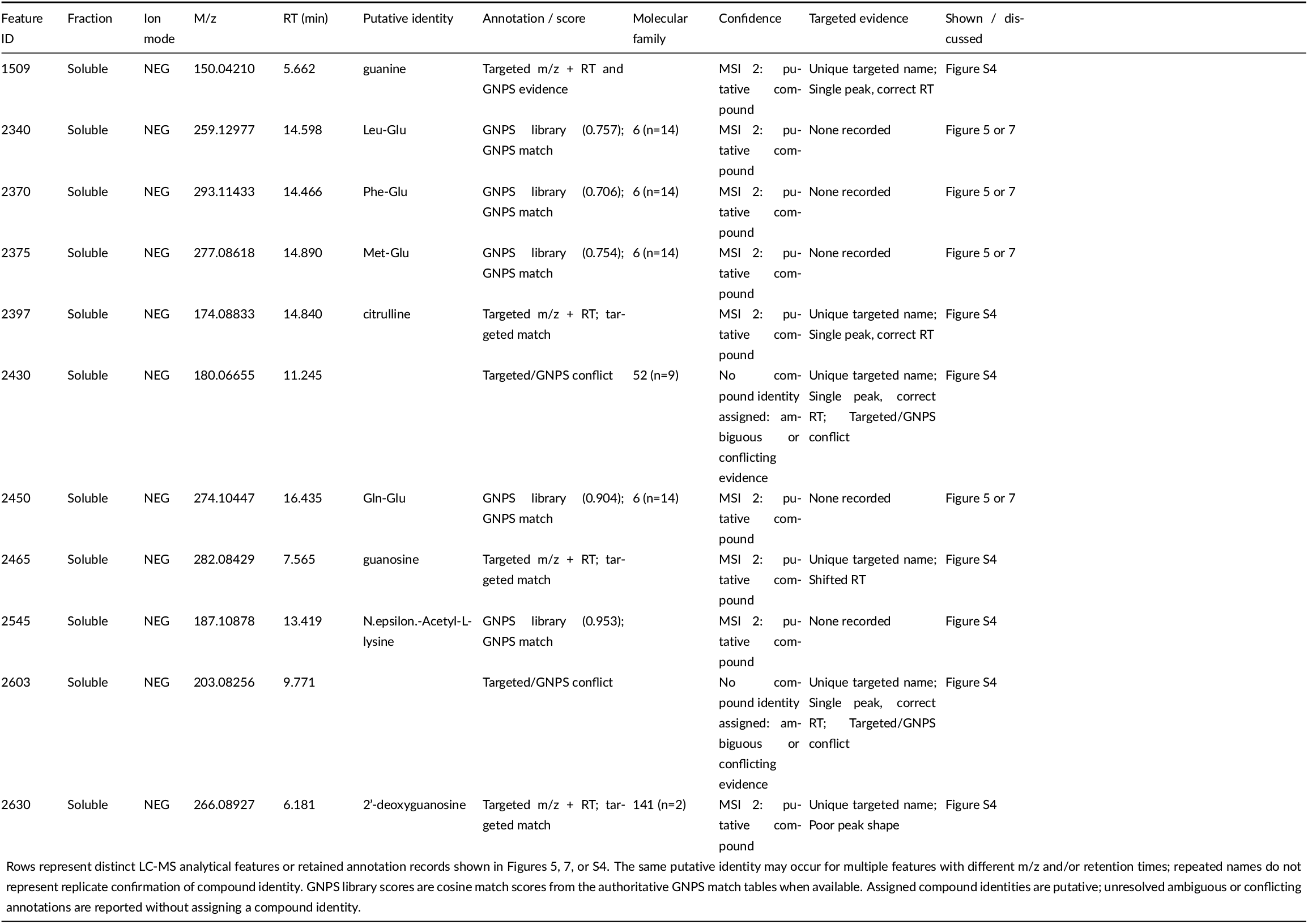
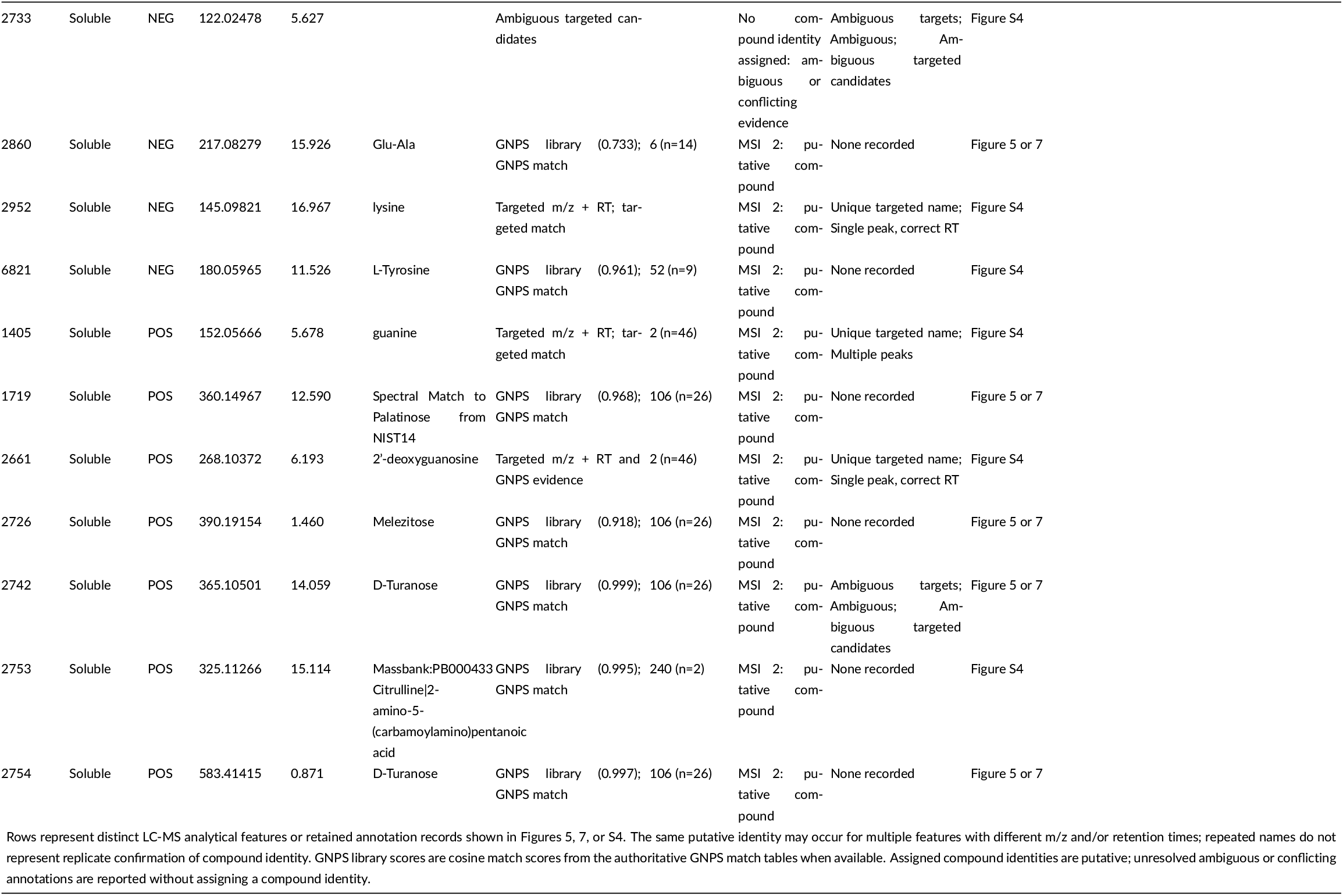
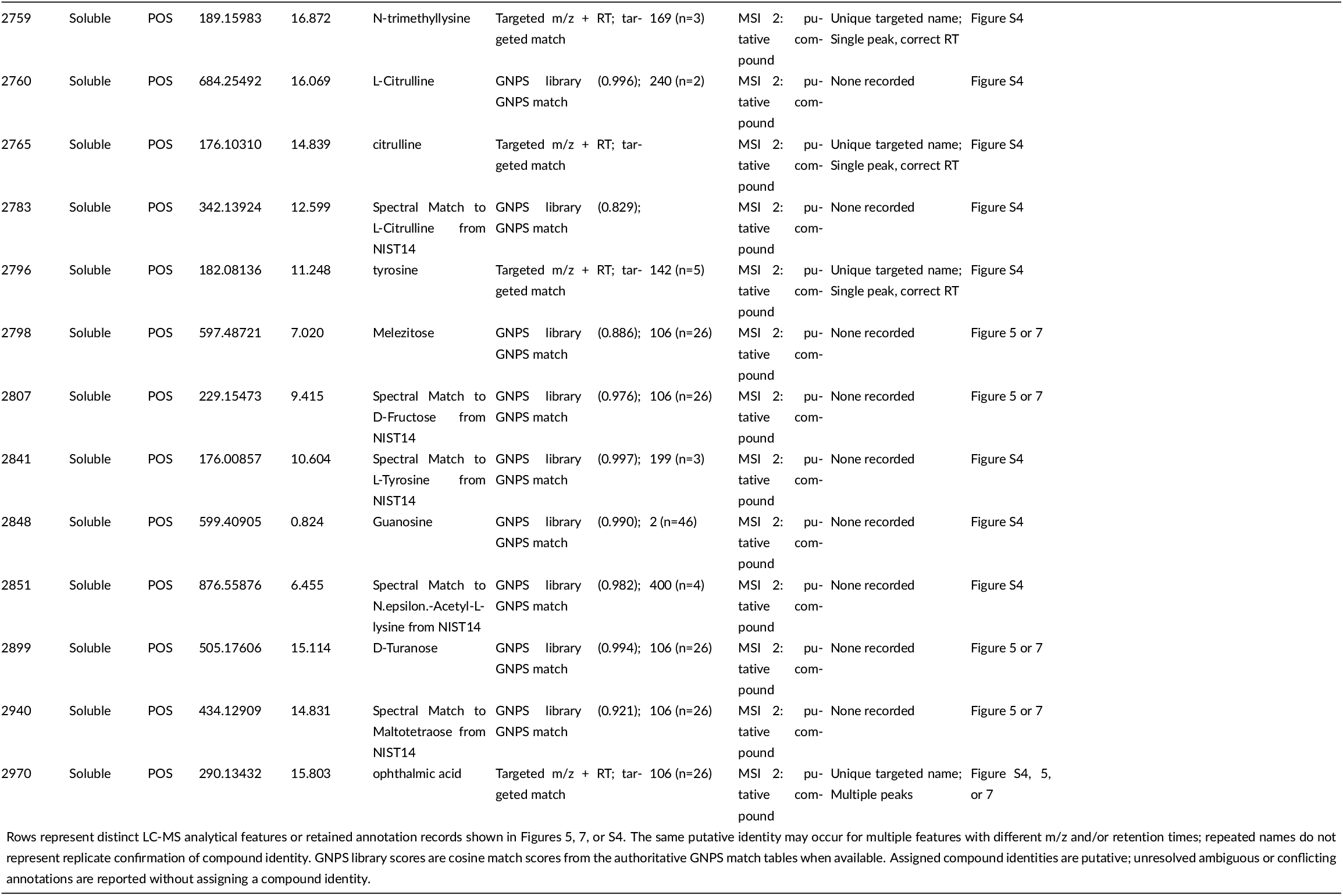
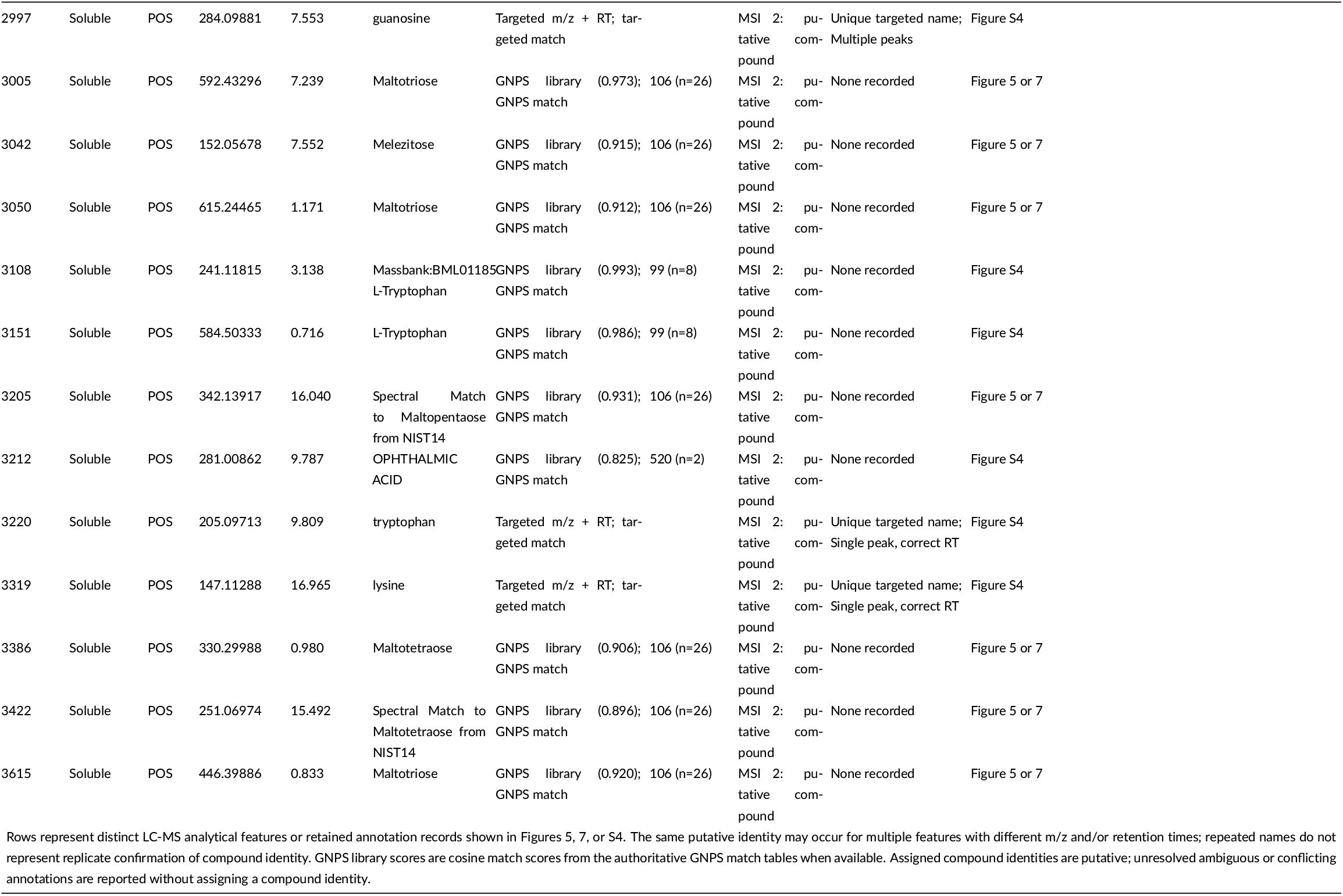
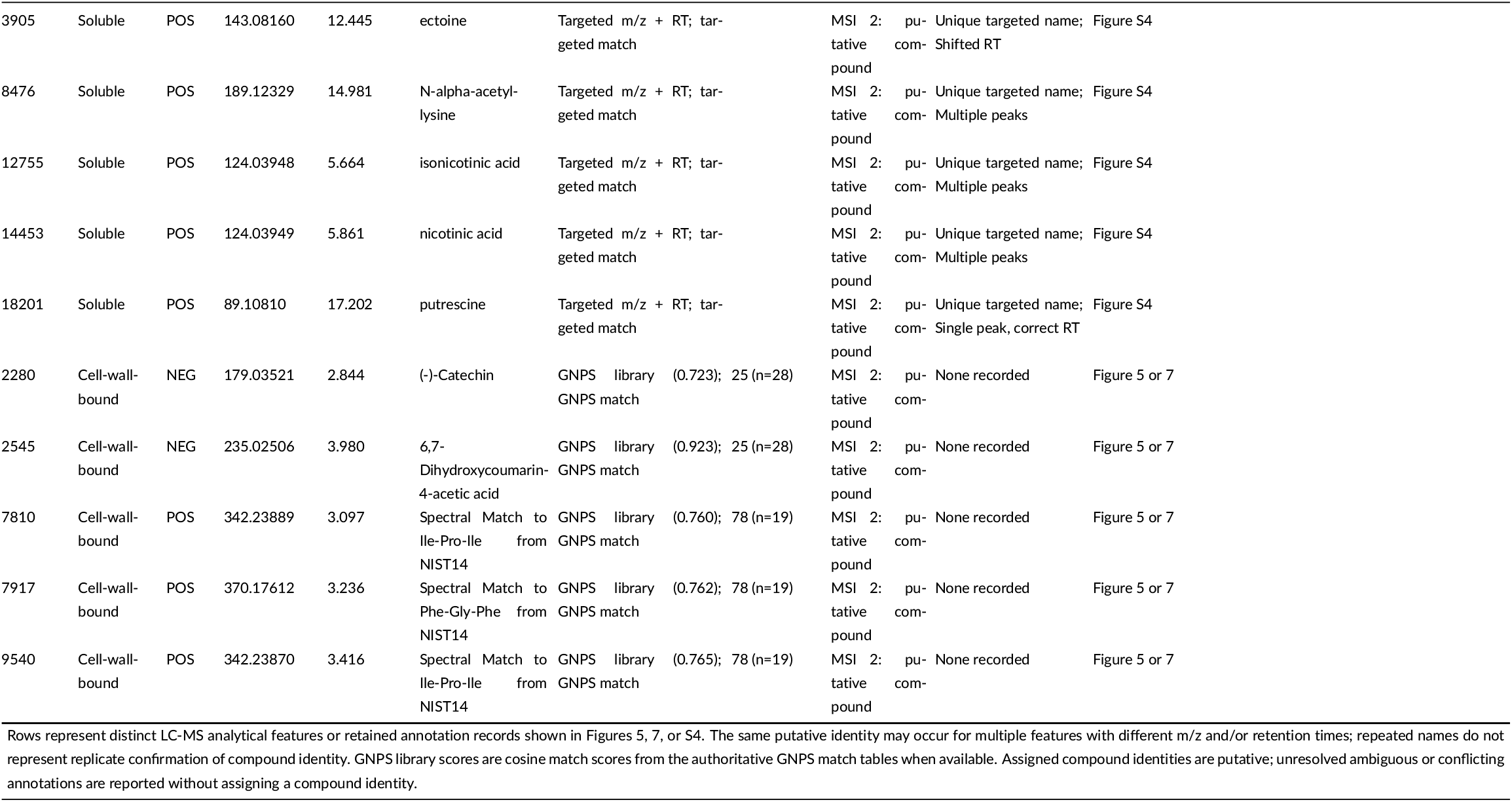
Metabolite annotation evidence relevant to displayed figures.

**TABLE S10.** Primary multivariate statistics and sensitivity summary.

| Dataset | Analysis | Term or groups | R<sup>2</sup> or F statistic | P | Adjusted P | Range or note |
| --- | --- | --- | --- | --- | --- | --- |
| SOL_HILIC_NEG | PERMANOVA | species | 0.22426975 | 1.000e-04 | 4.552e-04 | Primary full-data species-first model |
| SOL_HILIC_NEG | PERMANOVA | UVR | 0.079896568 | 1.000e-04 | 4.552e-04 | Primary full-data species-first model |
| SOL_HILIC_NEG | PERMANOVA | hydration | 0.30204207 | 1.000e-04 | 4.552e-04 | Primary full-data species-first model |
| SOL_HILIC_NEG | PERMANOVA | species:UVR | 0.043078866 | 0.0027000000 | 0.0075829787 | Primary full-data species-first model |
| SOL_HILIC_NEG | PERMANOVA | species:hydration | 0.047229969 | 0.0012000000 | 0.0041684211 | Primary full-data species-first model |
| SOL_HILIC_NEG | PERMANOVA | UVR:hydration | 0.016428608 | 0.17770000 | 0.23456400 | Primary full-data species-first model |
| SOL_HILIC_NEG | PERMANOVA | species:UVR:hydration | 0.016721338 | 0.16260000 | 0.22226667 | Primary full-data species-first model |
| SOL_HILIC_POS | PERMANOVA | species | 0.26473837 | 1.000e-04 | 4.552e-04 | Primary full-data species-first model |
| SOL_HILIC_POS | PERMANOVA | UVR | 0.083176063 | 1.000e-04 | 4.552e-04 | Primary full-data species-first model |
| SOL_HILIC_POS | PERMANOVA | hydration | 0.29139685 | 1.000e-04 | 4.552e-04 | Primary full-data species-first model |
| SOL_HILIC_POS | PERMANOVA | species:UVR | 0.044461928 | 0.0018000000 | 0.0055255814 | Primary full-data species-first model |
| SOL_HILIC_POS | PERMANOVA | species:hydration | 0.040977664 | 0.0026000000 | 0.0074608696 | Primary full-data species-first model |
| SOL_HILIC_POS | PERMANOVA | UVR:hydration | 0.014334928 | 0.19610000 | 0.24953077 | Primary full-data species-first model |
| SOL_HILIC_POS | PERMANOVA | species:UVR:hydration | 0.017073123 | 0.12190000 | 0.18330000 | Primary full-data species-first model |
| CWB_C18_NEG | PERMANOVA | species | 0.32517341 | 1.000e-04 | 4.552e-04 | Primary full-data species-first model |
| CWB_C18_NEG | PERMANOVA | UVR | 0.070224453 | 0.0069000000 | 0.015703448 | Primary full-data species-first model |
| CWB_C18_NEG | PERMANOVA | hydration | 0.037277813 | 0.093200000 | 0.16624865 | Primary full-data species-first model |
| CWB_C18_NEG | PERMANOVA | species:UVR | 0.025264486 | 0.25230000 | 0.30276000 | Primary full-data species-first model |
| CWB_C18_NEG | PERMANOVA | species:hydration | 0.028566384 | 0.18450000 | 0.24112871 | Primary full-data species-first model |
| CWB_C18_NEG | PERMANOVA | UVR:hydration | 0.031684257 | 0.13550000 | 0.19441304 | Primary full-data species-first model |
| CWB_C18_NEG | PERMANOVA | species:UVR:hydration | 0.017217448 | 0.51020000 | 0.52614375 | Primary full-data species-first model |
| CWB_C18_POS | PERMANOVA | species | 0.23537346 | 1.000e-04 | 4.552e-04 | Primary full-data species-first model |
| CWB_C18_POS | PERMANOVA | UVR | 0.094178785 | 2.000e-04 | 8.250e-04 | Primary full-data species-first model |
| CWB_C18_POS | PERMANOVA | hydration | 0.067420240 | 0.0035000000 | 0.0090588235 | Primary full-data species-first model |
| CWB_C18_POS | PERMANOVA | species:UVR | 0.033013545 | 0.11210000 | 0.18268148 | Primary full-data species-first model |
| CWB_C18_POS | PERMANOVA | species:hydration | 0.025246927 | 0.25820000 | 0.30451034 | Primary full-data species-first model |
| CWB_C18_POS | PERMANOVA | UVR:hydration | 0.044800013 | 0.035600000 | 0.069105882 | Primary full-data species-first model |
| CWB_C18_POS | PERMANOVA | species:UVR:hydration | 0.017080643 | 0.57800000 | 0.58719692 | Primary full-data species-first model |
| SOL_HILIC_NEG | dispersion | Within *S. ruralis*: four treatment cells | 0.18700466 | 0.90530000 | 0.90530000 | Primary dispersion test |
| SOL_HILIC_NEG | dispersion | Within *S. caninervis*: four treatment cells | 2.4550593 | 0.10630000 | 0.24803333 | Primary dispersion test |
| SOL_HILIC_NEG | dispersion | Eight species × UVR × hydration treatment cells | 1.1080000 | 0.39030000 | 0.54642000 | Primary dispersion test |
| SOL_HILIC_POS | dispersion | Within *S. ruralis*: four treatment cells | 0.19877106 | 0.89270000 | 0.90530000 | Primary dispersion test |
| SOL_HILIC_POS | dispersion | Within *S. caninervis*: four treatment cells | 1.7169950 | 0.20110000 | 0.35192500 | Primary dispersion test |
| SOL_HILIC_POS | dispersion | Eight species × UVR × hydration treatment cells | 0.84905347 | 0.55760000 | 0.68273333 | Primary dispersion test |
| CWB_C18_NEG | dispersion | Within *S. ruralis*: four treatment cells | 3.7612307 | 0.050100000 | 0.23380000 | Primary dispersion test |
| CWB_C18_NEG | dispersion | Within *S. caninervis*: four treatment cells | 0.69913211 | 0.58520000 | 0.68273333 | Primary dispersion test |
| CWB_C18_NEG | dispersion | Eight species × UVR × hydration treatment cells | 1.6180350 | 0.17260000 | 0.34520000 | Primary dispersion test |
| CWB_C18_POS | dispersion | Within *S. ruralis*: four treatment cells | 3.3797746 | 0.036200000 | 0.23380000 | Primary dispersion test |
| CWB_C18_POS | dispersion | Within *S. caninervis*: four treatment cells | 1.4318747 | 0.28630000 | 0.44535556 | Primary dispersion test |
| CWB_C18_POS | dispersion | Eight species × UVR × hydration treatment cells | 2.0082679 | 0.088600000 | 0.24803333 | Primary dispersion test |
| CWB_C18_NEG | term-order sensitivity | UVR |  |  |  | R<sup>2</sup> 0.06375–0.1474; P 0.0057–0.0484 |
| CWB_C18_NEG | term-order sensitivity | UVR:hydration |  |  |  | R<sup>2</sup> 0.02957–0.09609; P 0.1148–0.7431 |

**TABLE S10** Primary multivariate statistics and sensitivity summary
| Dataset | Analysis | Term or groups | R<sup>2</sup> or F statistic | P | Adjusted P | Range or note |
| --- | --- | --- | --- | --- | --- | --- |
| CWB_C18_NEG | term-order sensitivity | hydration |  |  |  | R2 0.0347–0.1086; P 0.0932–0.166 |
| CWB_C18_NEG | term-order sensitivity | species |  |  |  | R2 0.254–0.3256; P 1e-04–1e-04 |
| CWB_C18_NEG | term-order sensitivity | species:UVR |  |  |  | R2 0.02503–0.0269; P 0.2471–0.2671 |
| CWB_C18_NEG | term-order sensitivity | species:UVR:hydration |  |  |  | R2 0.01722–0.02451; P 0.3301–0.5102 |
| CWB_C18_NEG | term-order sensitivity | species:hydration |  |  |  | R2 0.02194–0.02857; P 0.1845–0.4347 |
| CWB_C18_POS | term-order sensitivity | UVR |  |  |  | R2 0.08417–0.1713; P 1e-04–0.003 |
| CWB_C18_POS | term-order sensitivity | UVR:hydration |  |  |  | R2 0.04035–0.0932; P 0.031–0.3598 |
| CWB_C18_POS | term-order sensitivity | hydration |  |  |  | R2 0.0613–0.1272; P 0.0035–0.0252 |
| CWB_C18_POS | term-order sensitivity | species |  |  |  | R2 0.1888–0.2361; P 1e-04–1e-04 |
| CWB_C18_POS | term-order sensitivity | species:UVR |  |  |  | R2 0.0323–0.0343; P 0.1043–0.1336 |
| CWB_C18_POS | term-order sensitivity | species:UVR:hydration |  |  |  | R2 0.01708–0.02195; P 0.4503–0.5783 |
| CWB_C18_POS | term-order sensitivity | species:hydration |  |  |  | R2 0.02267–0.02525; P 0.2582–0.4099 |
| SOL_HILIC_NEG | term-order sensitivity | UVR |  |  |  | R2 0.0799–0.164; P 1e-04–0.0054 |
| SOL_HILIC_NEG | term-order sensitivity | UVR:hydration |  |  |  | R2 0.01643–0.04488; P 0.1638–0.231 |
| SOL_HILIC_NEG | term-order sensitivity | hydration |  |  |  | R2 0.302–0.4945; P 1e-04–1e-04 |
| SOL_HILIC_NEG | term-order sensitivity | species |  |  |  | R2 0.2243–0.2243; P 1e-04–1e-04 |
| SOL_HILIC_NEG | term-order sensitivity | species:UVR |  |  |  | R2 0.04308–0.04308; P 0.0027–0.003 |
| SOL_HILIC_NEG | term-order sensitivity | species:UVR:hydration |  |  |  | R2 0.01672–0.01672; P 0.1626–0.1667 |
| SOL_HILIC_NEG | term-order sensitivity | species:hydration |  |  |  | R2 0.04723–0.04723; P 0.0012–0.0026 |
| SOL_HILIC_POS | term-order sensitivity | UVR |  |  |  | R2 0.08318–0.1765; P 1e-04–0.0014 |
| SOL_HILIC_POS | term-order sensitivity | UVR:hydration |  |  |  | R2 0.01433–0.0474; P 0.1199–0.2668 |
| SOL_HILIC_POS | term-order sensitivity | hydration |  |  |  | R2 0.2914–0.486; P 1e-04–1e-04 |
| SOL_HILIC_POS | term-order sensitivity | species |  |  |  | R2 0.2647–0.2647; P 1e-04–1e-04 |
| SOL_HILIC_POS | term-order sensitivity | species:UVR |  |  |  | R2 0.04446–0.04446; P 0.001–0.0018 |
| SOL_HILIC_POS | term-order sensitivity | species:UVR:hydration |  |  |  | R2 0.01707–0.01707; P 0.1219–0.1251 |
| SOL_HILIC_POS | term-order sensitivity | species:hydration |  |  |  | R2 0.04098–0.04098; P 0.0022–0.0026 |

The CSV contains complete PERMANOVA, dispersion, and term-order sensitivity records, including sample-52-inclusion models. The PDF reports only the primary full-data species-first PERMANOVA, primary dispersion tests, and term-order sensitivity ranges; sample-52-inclusion rows remain identifiable in the CSV. PERMANOVA and dispersion permutation P values were adjusted separately by Benjamini–Hochberg across the complete PERMANOVA and dispersion result tables, respectively.

**TABLE S11.** Robustness and sensitivity analysis summary.

| Sensitivity analysis | Dataset | Median Jaccard | Minimum Jaccard |
| --- | --- | --- | --- |
| Sample 52 inclusion | Cell-wall-bound, negative | 0.60 | 0.00 |
| Sample 52 inclusion | Cell-wall-bound, positive | 0.71 | 0.46 |
| Linear injection-order adjustment | Cell-wall-bound, negative | 0.76 | 0.76 |
| Linear injection-order adjustment | Cell-wall-bound, positive | 0.71 | 0.71 |
| Linear injection-order adjustment | Soluble, negative | 0.99 | 0.99 |
| Linear injection-order adjustment | Soluble, positive | 0.99 | 0.99 |
| Alternative normalization | Cell-wall-bound, negative | 0.89 | 0.00 |
| Alternative normalization | Cell-wall-bound, positive | 0.73 | 0.25 |
| Alternative normalization | Soluble, negative | 0.89 | 0.75 |
| Alternative normalization | Soluble, positive | 0.92 | 0.74 |
| Control-associated-feature threshold | Cell-wall-bound, negative | 0.94 | 0.69 |
| Control-associated-feature threshold | Cell-wall-bound, positive | 0.93 | 0.00 |
| Control-associated-feature threshold | Soluble, negative | 0.99 | 0.94 |
| Control-associated-feature threshold | Soluble, positive | 0.99 | 0.89 |
| Contaminant-family exclusion | Cell-wall-bound, negative | 1.00 | 1.00 |
| Contaminant-family exclusion | Cell-wall-bound, positive | 0.98 | 0.33 |
| Contaminant-family exclusion | Soluble, negative | 1.00 | 1.00 |
| Contaminant-family exclusion | Soluble, positive | 1.00 | 1.00 |

Jaccard similarity summarizes overlap in significant-feature sets between each sensitivity analysis and the primary analysis across applicable contrasts and model terms. The PDF reports the finite Jaccard summaries for linear injection-order adjustment; complete contrast-level sensitivity results and injection-order diagnostic records are provided in the accompanying Table S11.csv. Jaccard values are descriptive; no binary robustness threshold was imposed.

